# WiChR, a highly potassium selective channelrhodopsin for low-light two-photon neuronal inhibition

**DOI:** 10.1101/2022.07.02.498568

**Authors:** Johannes Vierock, Enrico Peter, Christiane Grimm, Andrey Rozenberg, Alejandro G. Castro Scalise, Sandra Augustin, Dimitrii Tanese, Benoît C. Forget, Valentina Emiliani, Oded Béjà, Peter Hegemann

**Affiliations:** Institut für Biologie, Experimentelle Biophysik, Humboldt-Universität zu Berlin, Invalidenstrasse 42, 10115 Berlin, Germany; Neuroscience Research Center, Charité - Universitätsmedizin Berlin, Berlin Germany; Wavefront Engineering Microscopy Group, Photonics Department, Institut de la Vision, Sorbonne Université, INSERM, CNRS, Paris, France; Faculty of Biology, Technion – Israel Institute of Technology, Haifa 32000, Israel

**Author notes:** These authors contributed equally to this work: Johannes Vierock, Enrico Peter, Christiane Grimm. Corresponding authors (J.V.); (P.H.).

## Abstract

The electric excitability of muscle, heart and brain tissue relies on the precise interplay of Na^+^- and K^+^-selective ion channels. The involved ion fluxes are controlled in optogenetic studies using light-gated channelrhodopsins (ChRs). While non-selective cation-conducting ChRs are well-established for excitation, K^+^-selective ChRs (KCRs) for efficient inhibition have only recently come into reach. Here, we report the molecular analysis of recently discovered KCRs from the stramenopile *Hyphochytrium catenoides* and identify a novel type of hydrophobic K^+^-selectivity filter. Next, we demonstrate that the KCR signature motif is conserved in related stramenopile ChRs. Among them, WiChR from *Wobblia lunata* features an unmatched 80-fold preference for K^+^ over Na^+^, stable photocurrents under continuous illumination and a prolonged open state lifetime. Well expressed in neurons, WiChR allows two-photon inhibition at low irradiance and reduced tissue heating,_recommending WiChR as the long-awaited efficient and versatile optogenetic inhibitor.

## Introduction

Light-gated ion channels are used in modern life sciences from plant physiology to systems neuroscience to control the electrical activity of cells with the temporal and spatial precision of light (*1*). Simultaneous illumination of individual cells within larger brain areas is reliably achieved by holographic light shaping and two photon (2P) excitation (*2*). The employed light switches are with few exceptions channelrhodopsins (ChRs) that absorb light via a retinal chromophore bound to the seven-transmembrane-helix protein as a retinal Schiff base. First described as the photoreceptor responsible for phototaxis in the green alga *Chlamydomonas reinhardtii* (*3*), ChRs have been discovered across different families of microalgae with more than 800 described representatives today (*4, 5*).

While cation-conducting channelrhodopsins (CCRs) are well established for excitation, neuronal inhibition is most frequently achieved with anion-conducting ChRs (ACRs). However, variability of internal Cl^-^ concentration, which is high in axonal and synaptic terminals, in cardiomyocytes and during development, may cause depolarization instead of hyperpolarization and limits the application of ACRs (*6–8*).Other inhibitory tools such as light-driven pumps and inhibitory G_i/o_-coupled Opto-GPCRs also have significant limitations. Optogenetic pumps require high expression levels and continuous bright illumination due to a strict one photon/one charge ratio and can have undesired side effects (*9*). Opto-GPCRs require cell type–specific examination of engaged signaling cascades and might activate multiple signaling pathways with limited temporal on-off control (*10*).

An attractive alternative for optogenetic inhibition and a long-pursued tool is a potassium selective CCR that would hyperpolarize the neuronal membrane despite high extracellular Na^+^ concentration and mimic the major endogenous repolarization processes based on K^+^ efflux. Engineering a K^+^-selective ChR is an ambitious goal considering major structural differences between ChRs and natural K^+^ channels. Whereas ChRs conduct protons and partly hydrated ions along a sequence of water filled cavities through an overall asymmetric and still structurally unknown pore within the rhodopsin monomer itself (*11*), K^+^ channels feature a highly conserved and symmetric selectivity filter with four subsequent K^+^ binding sites for K^+^ dehydration in the symmetry axis of the tetrameric channel (*12*). Accordingly, first-generation K^+^ channels combined light sensitive modules such as UV-switchable azobenzene compounds, blue-light sensitive LOV-domains or blue-light activated cyclases with ligand-, viral and cAMP-gated K^+^ channels as single or two component systems (*13–15*). The need for an additional co-factor, low expression, slow kinetics, a limited dynamic range and putative side effects limited their application in neuroscience to date. Structure-guided molecular engineering of ChRs led only to minor improvements in the *P*_K+_/*P*_Na+_ permeability ratio (*11*), whereas modifications of the extracellular release channel of the light-driven Na^+^-pump KR2 of *Dokdonia eikasta* surprisingly generated an operational light-activated K^+^ channel with substantial potassium conductance, but only at alkaline pH (*16*).

A veritable breakthrough represents the recent discovery of two natural potassium-conducting channelrhodopsins (KCRs) in the stramenopile *Hyphochytrium catenoides* (HcKCR1 and HcKCR2) that were K^+^-selective enough to inhibit action potential firing in neuronal slices (*17*). Both KCRs have a bacteriorhodopsin-like DTD motif in membrane helix 3, a feature shared with the related cryptophyte CCRs that were first discovered in *Guillardia theta* (*18*)and include the green-light activated non-selective cation channel ChRmine from *Rhodomonas lens*, the structure of which was recently solved by Cryo-EM (*19, 20*). Despite an extensive biophysical characterization by Govorunova and colleagues, the molecular determinants of K^+^-selectivity in these newly identified KCRs remain unknown.

Here, we inspect the potassium selectivity of both HcKCRs, quantify their remaining Na^+^-conductance and identify key residues for K^+^-conductance and selection over Na^+^. A search for new CCRs containing the identified K^+^-selectivity signature motif, revealed WiChR from *Wobblia lunata* with improved K^+^-selectivity, negligible photocurrent inactivation and improved light sensitivity compared to both previously characterized KCRs. Expressed in hippocampal brain slices, WiChR showed reduced depolarization at different membrane potentials and prolonged inhibition with minimally invasive illumination protocols. Highly K^+^-selective, well expressed in mammalian cells and sensitive to holographic two-photon excitation, we recommend WiChR as a new tool for multi-target single cell inhibition in large neuronal networks.

## Results

### HcKCRs are K^+^-selective channels with substantial Na^+^-conductance

We expressed both HcKCRs in ND7/23 cells and compared photocurrents at different ionic conditions with those of ChRmine (Fig. 1B). In the presence of high intra- and extracellular K^+^, large photocurrents confirmed efficient potassium conductance for all three channels with even higher photocurrent densities for both HcKCRs than for ChRmine. Extracellular buffer exchange had no effect on photocurrents of ChRmine that conducts both cations equally well, but caused a significant reduction of inward currents and a strong shift in the reversal potential for both HcKCRs confirming their pronounced preference for K^+^ over Na^+^ (Fig. 1C-F). Further, exchange of Na^+^ to the larger cation NMG^+^ abolished all remaining inward currents in HcKCRs and shifted the reversal potential in all three channels indicating that in ChRmine, as well as in HcKCRs Na^+^ ions are still conducted to an considerable extent. Moreover, at voltages close to *E*_rev_ extended illumination led to a directional change of the HcKCR1 photocurrents and to a transition from a K^+^ outward current to a Na^+^-influx, revealing an increasing Na^+^-selectivity over time. The concomitant *E*_rev_ shifted by about 10 mV and reflects a permeability ratio (*P*_K+_/*P*_Na+_) drop from 29 ± 4 to 18 ± 2 (Fig. S1). At the same time, HcKCR1 photocurrents inactivate by 30 to 60 % depending on the voltage and ionic conditions and recover biphasically within seconds in the dark (Fig. S2).

**Fig. 1.**
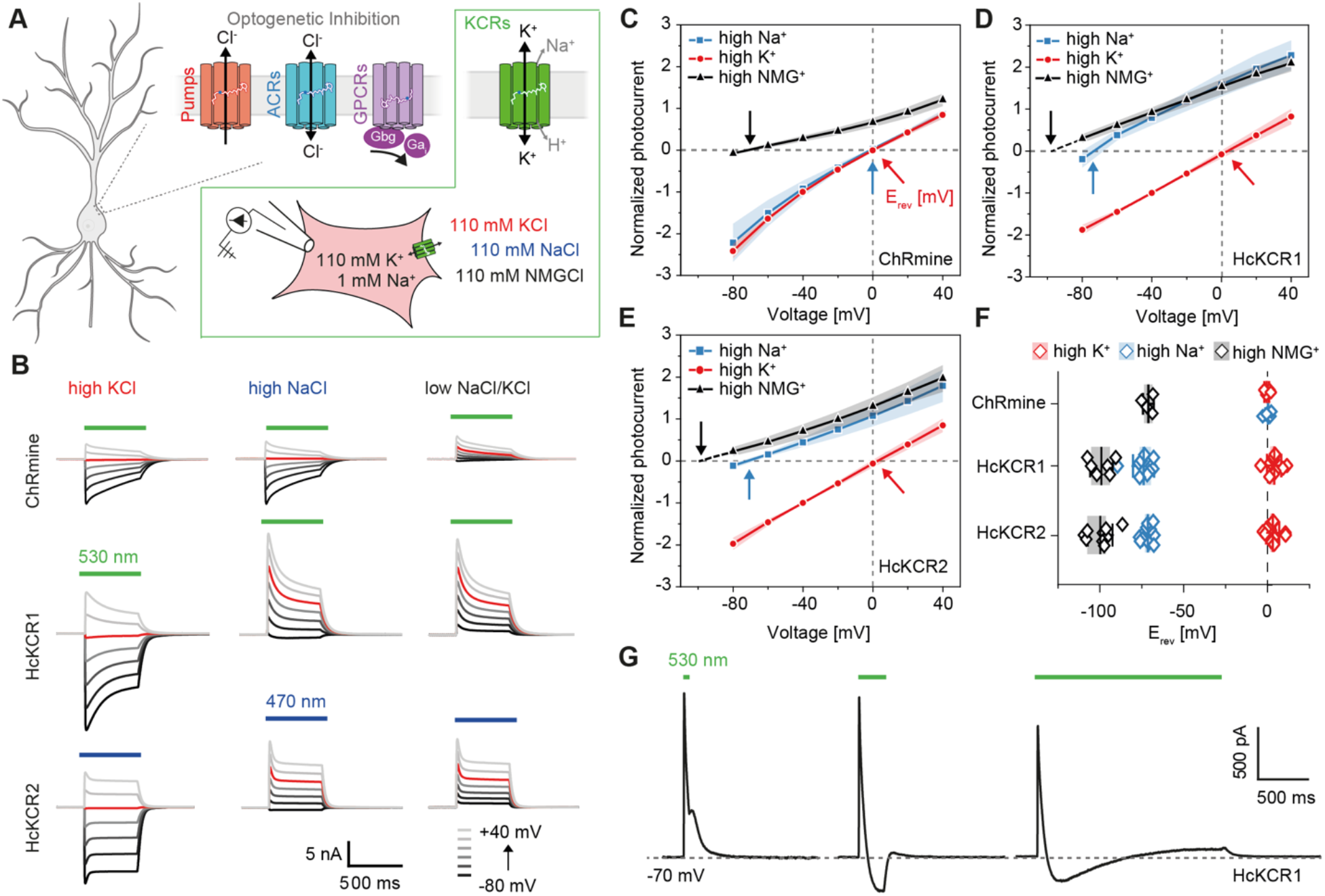
K^+^-selectivity of HcKCRs. (**A**) Established optogenetic inhibitory tools including light driven ion pumps (red), anion conducting channelrhodopsins (blue), and G_i/q_-coupled opsins (purple). (**B**) Photocurrents of ChRmine, HcKCR1 and HcKCR2 at 110 mM K^+^_e_, Na^+^_e_, or NMG^+^_e_. and 110 mM K^+^_i_ gluconat. **C**-**E** I(E) relations of the normalized peak photocurrent amplitudes (Mean ± SD and n = 7/7/4 for ChRmine, n=10/10/6 for HcKCR1 and n=13/13/9 for HcKCR2) (**F**) E_rev_ values for C - E. (**G**) Photocurrents of HcKCR1 at −70 mV at high Na^+^_e_ upon 50 ms, 250 ms and 2 s illumination.

### K^+^-selectivity of HcKCRs relies on aromatic residues in the extracellular half pore

In the next set of experiments, we identified key amino acids relevant for efficient K^+-^conductance and selectivity. All known ChRs conduct ions along a sequence of preformed water-filled cavities between transmembrane helices 1, 2, 3 and 7 (*11*). The putative HcKCR1 pore is spanning from the extracellular D87 in transmembrane helix 2 to the intracellular D116 in helix 3 (Fig. 2A). The aspartates D105 and D229 in the center of the pore constitute the counterion complex that stabilizes the protonated retinal Schiff base charge. The extracellular half pore is lined by a set of aromatic residues including Y81, F88, W210, Y222 and an unusual tryptophan W102 at a position most frequently occupied by arginine in other microbial rhodopsins.

**Fig. 2.**
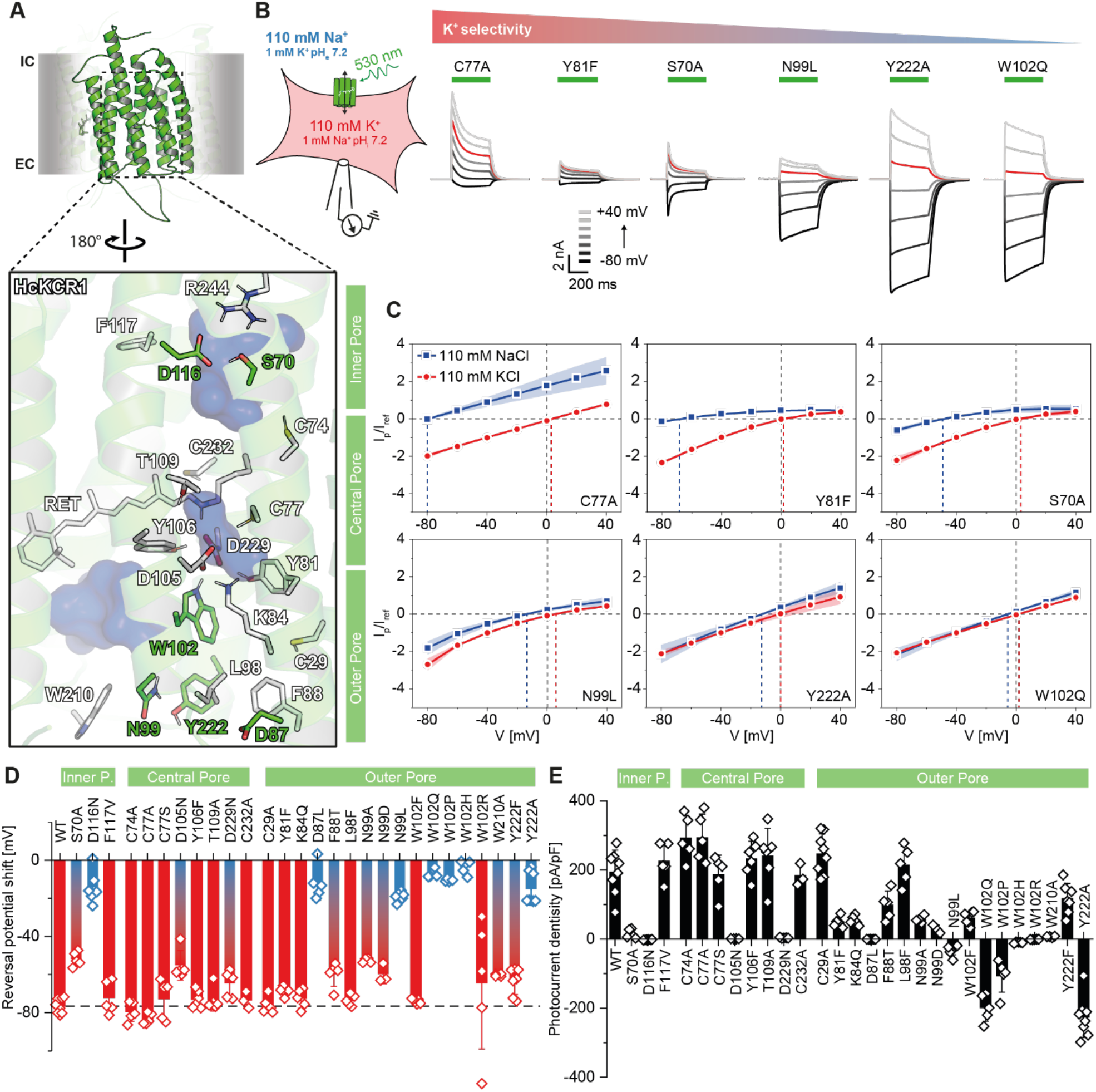
Molecular determinants for the K^+^-conductance in KCRs. (**A**) Equilibrated AlphaFold2 3D model of HcKCR1 with important residues lining the putative pore. (**B**) Representative photocurrents of pore lining HcKCR1 mutants at 110 mM K^+^_i_ and 110 mM Na^+^_e_. (**C**) I(E) relations of the normalized peak-photocurrent amplitudes of selected mutants. (**D**)ΔE_rev_ for extracellular buffer exchange from high K^+^_e_ to Na^+^_e_ (**E**) Photocurrent density at - 40 mV and 110 mM Na^+^_e_. All plots show (Mean ± SD, n = 3-8)

Mutations of most of the pore-lining residues had little impact on the K^+^-conductance and selectivity (Fig. S3+S4). Accordingly, photocurrents of the central gate mutant C77A still resembled WT currents (Fig. 2B-C), whereas mutations of F81, F88 or S70 that are located in close contact with the pore aspartates caused strong reduction of outward currents and impaired K^+^-selection as indicated by the diminished Δ*E*_rev_. A nearly complete loss of K^+^-selectivity was observed for the outer pore mutant N99L and for substitutions of W102 and Y222 by non-aromatic residues such as W102Q and Y222A. For all three mutants, inward currents were large at both high extracellular Na^+^ or K^+^ (Fig. 2C) with little reversal potential shift between the two conditions (Fig. 2D), similar to the non-selective cation channel ChRmine (Fig. 1C+F). K^+^-selectivity was also compromised in D87 and D116 mutants but came with a near-complete loss of stationary photocurrents (Fig. 2E). Mutations of the counterion complex also showed reduced photocurrent amplitudes but, by contrast, largely preserved K^+^-selectivity. In conclusion, all four aspartates of the pore and their interaction partners S70 and Y81 are essential for channel gating and proper pore formation in KCRs, whereas W102 and Y222 constitute essential elements of a unique hydrophobic K^+^-selectivity filter.

Based on our mutational analysis, we define the quintet D87, N99, W102, D116 and Y222 as the KCR signature motif that requires functional support by S70, F88 and D105 as important interaction partners for efficient K^+^-conductance. (Fig. 3A).

**Fig. 3.**
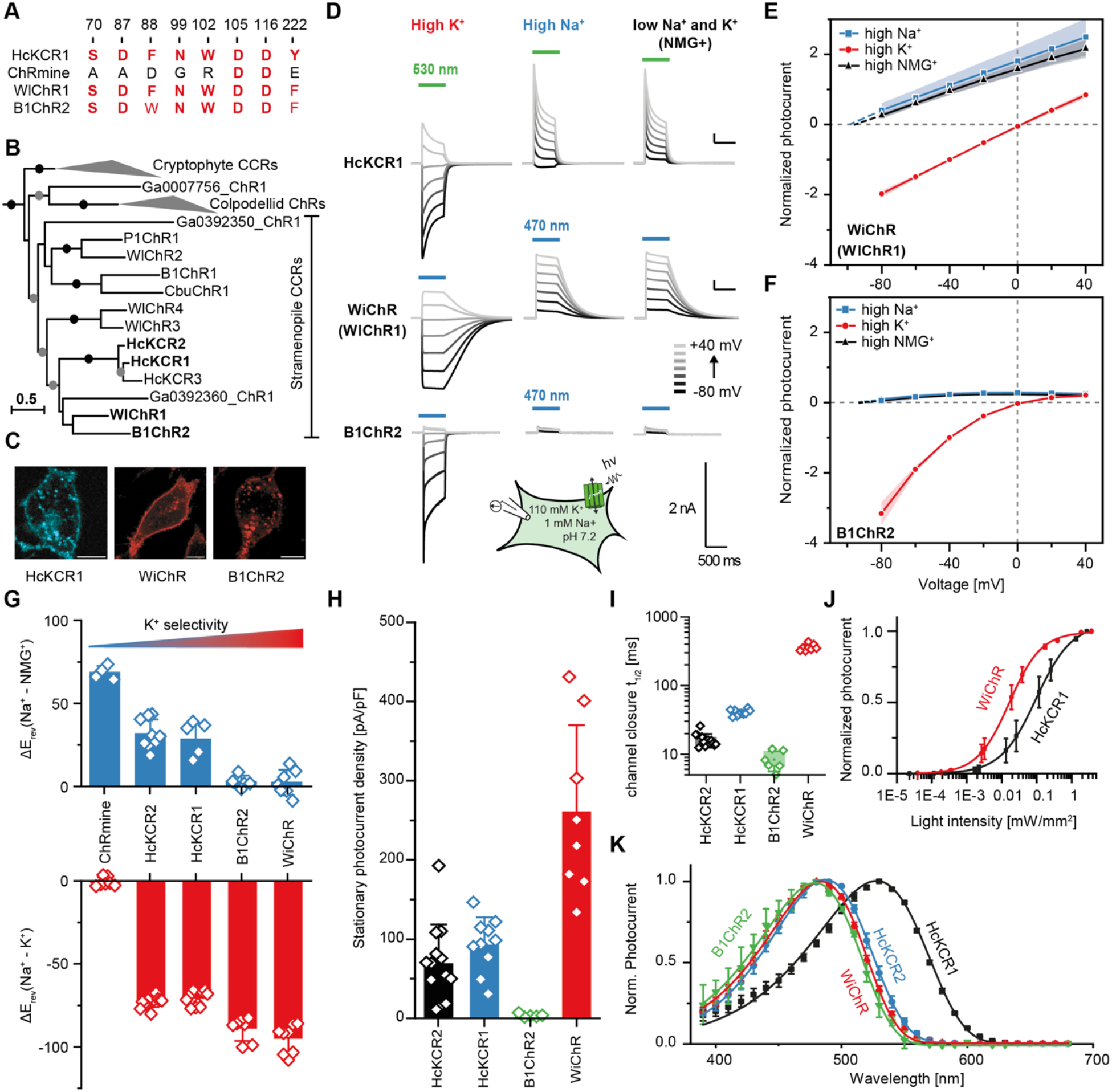
Identification and characterization of WiChR. (**A).** The 8 amino acid motif of the K^+^-selectivity filter in red (see also Fig S6A and the complete alignment in Fig. S8 and Suppl. Data File S4). (**B**) Phylogenetic relationships among the stramenopile CCRs (see the complete phylogenetic tree of the channelrhodopsin family in Fig. S6A and Suppl. Data File S2). Dots represent ultra-fast bootstrap support values: ≥ 80 (gray) and ≥ 95 (black). (**C**)Cellular distribution of the fluorescence tagged KCRs. (**D**) Representative photocurrents of HcKCR1, WiChR, and B1ChR2 at high K^+^_e_, Na^+^_e_, or NMG^+^_e_. and intracellular 110 mM K^+^_i_ gluconate. (**E) & (F**) I(E) relations of the peak photocurrent amplitudes of WiChR/WlChR1 (n = 8/8/7) and B1ChR2 (n = 7/7/5). (**G**) ΔE_rev_ of stationary photocurrents (after 500 ms light) for extracellular buffer exchange from high K^+^_e_ to Na^+^_e_ (bottom) and high Na^+^_e_ to NMG^+^_e_. (top) (**H**) stationary photocurrent densities with high Na^+^e and −40 mV. (**I**)photocurrent closure after 500 ms recordings. (**J**)light titration of the WiChR (n = 5) and HcKCR1 (n=5) peak photocurrents. (**K**)Action spectra of peak photocurrents (n= 6 for all four channels). All plots show (mean ± SD).

### Identification of new candidate KCRs

A systematic search revealed that KCRs from *H. catenoides* (HcKCR1 and HcKCR2) are part of a bigger clade that unites ChRs from a small set of stramenopile protists and related metatranscriptomic sequences (Fig. 3B, Fig. S6 and S7A). The four non-hyphochytridiomycete stramenopiles that encode members of this ChR group are heterotrophic flagellates from the clade of Opalozoa: placidids *Wobblia lunata* and Placidida sp. Caron Lab Isolate and anoecids *Cafeteria burkhardae* and *Bilabrum* sp. (the latter identified using molecular markers). Together with cryptophyte (B)CCRs (including ChRmine) and a novel clade of uncharacterized ChRs from colpodellid alveolates these proteins form a well-supported monophylum (Fig. 3B, Fig. S6A). Most stramenopile CCRs share the functionally important DTD motif of bacteriorhodopsin (D85, T89, D96) in transmembrane helix 3 with cryptophyte BCCRs – and all of the minor variations of the motif (DSD, DSE, ETD and DLD) preserve both of the carboxylates - whereas in related colpodellid proteins only the second aspartate is conserved (GGD motif) (Fig. S6B). Clustering of the stramenopile CCRs and cryptophyte (B)CCRs indicates that cation conductivity is ancestral in this lineage.

Among the collected stramenopile ChRs from this clade, two proteins, WlChR1 from *W. lunata* and B1ChR2 from *Bilabrum* sp. contained a nearly complete KCR signature motif with only one conservative substitution at the position of Y222 for a phenylalanine - a substitution that preserved K^+^-selectivity in HcKCR1 in our mutational analysis (Fig. S4).

### Electrophysiological characterization of WlChR1 (WiChR) in cultured cells

Expressed in ND7/23 cells, WlChR1 showed excellent membrane targeting (Fig. 3C) and large photocurrents with almost no inactivation during prolonged illumination (Fig. 3D). In contrast, B1ChR2 aggregated substantially in intracellular compartments and produced photocurrents with rapid inactivation and strong inward rectification. Photocurrent recordings at different K^+^_e_ concentrations confirmed a high K^+^-selectivity for both channels. Strikingly, WlChR1 as well as B1ChR2 even showed a complete loss of inward directed currents upon exchange of extracellular K^+^ to Na^+^ (Fig. 3E +F). The corresponding larger ΔE_rev_ for WlChR1 compared to HcKCR1 (Fig. 3G) is interpreted on the basis of the Goldmann-Hodgkin-Katz voltage equation as a remarkable increase in relative K^+-^conductance from *P*_K+_/*P*_Na+_ = 20 ± 4 for HcKCR1 to *P*_K+_/*P*_Na+_ = 80 ± 40 for WlChR1. Improved K^+^-selectivity is accompanied by enlarged stationary photocurrents (Fig. 3H) - explained by the good expression and reduced photocurrent inactivation - and slow channel closure (Fig. 3I) that both result in an improved operational light sensitivity (Fig. 3J). The blue-shifted action spectrum compared to HcKCR1 favors combination with orange or red absorbing actuators and sensors and largely overlaps with HcKCR2 (Fig. 3K).

Considering WlChR1 as a highly promising tool for optogenetic inhibition, we term it WiChR for *<u>W</u>obblia* <u>i</u>nhibitory <u>Ch</u>annel<u>R</u>hodopsin.

### 2P holographic inhibition of neurons in organotypic slices

To explore neuronal application, we expressed HcKCR1 and WiChR in organotypic slices from the mouse hippocampus and compared their inhibitory performance using holographic 2-photon patterned stimulation and a low repetition rate pulsed laser at 1030 nm (Fig. 4A). Both KCRs expressed well (Fig. 4B) and showed no signs of toxicity (Fig. S12). We injected depolarizing square currents (1 s) to induce action potential firing and illuminated in the middle of the current injection. Both HcKCR1 and WiChR completely inhibited spiking at current injections well above the respective rheobase of the neuron (Fig. 4B). While for HcKCR1 continuous illumination was needed, for WiChR a single 5 ms pulse was sufficient to achieve complete cessation of firing for the entire 500 ms (Fig. 4C). Since extended periods of inhibition are often needed during in vivo applications, pulsed illumination can reduce the local heating in the tissue compared to continuous illumination. Hence, we compared inhibition with continuous illumination to the application of 5 ms pulses at 10, 5 and 2 Hz and found that WiChR achieves complete cessation of action potential firing down to a frequency of 2 Hz while HcKCR1 needs continuous illumination for the same degree of inhibition (Fig. S13A,B). Next, we simulated the local temperature rise (*21*)for the holographic spot (12 μm diameter, 31 μW/μm^2^) and compared continuous with pulsed illumination. We found that heat is accumulating with continuous illumination, but the pulsed protocol at 10, 5 or 2 Hz allows dissipation of the heat before the next pulse (Fig. S13C). While for one holographic spot the local temperature rise of 0.23 K (continuous) and 0.14 K (pulsed) is still tolerable, most applications will demand inhibition of more than one neuron in a given region. Here, our simulations show that with a 5 ms illumination pulsed at 2 Hz, peak heating stays below 1 K and no heat is accumulated over time even when placing up to 40 holographic spots within a small 100 x 100 μm^2^ area. On the contrary, for continuous illumination, heat is accumulated and increases linearly with the number of spots reaching almost 4 K for the same 40 spots (Fig. S13D).

**Fig. 4:**
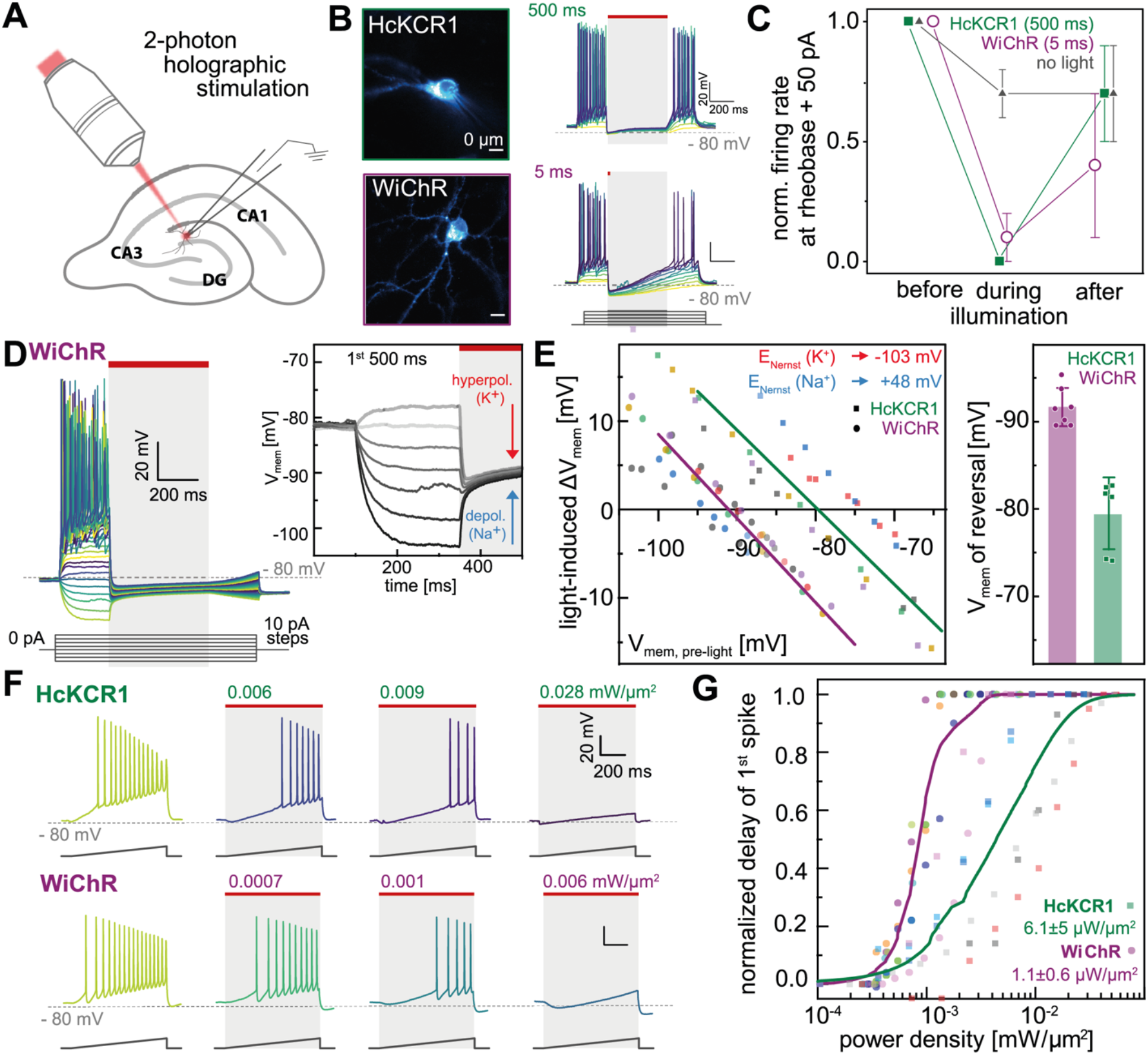
Neuronal inhibition with 2-photon holographic stimulation of WiChR in organotypic hippocampal slices. (**A**) Whole-cell patch clamp recordings of HcKCR1 and WiChR with 2P holographic illumination (1030 nm); liquid junction potential corrected for all recordings (**B**) Epifluorescence images and recordings of square depolarizing current injections (1 s) with a 500 ms period of photoinhibition within this time; continuous illumination for HcKCR1 and 5 ms for WiChR (29 μW/μm^2^) (**C**) Normalized firing rate 50 pA above rheobase for HcKCR1, WiChR (as in B) and without light, mean±SD, n=6-8 each. (**D**) Like (A) with hyperpolarizing current injections (**E**) Extrapolation of membrane potential where illumination induces depolarization instead of hyperpolarization; points individual recordings and lines average. (**F**) Depolarizing somatic ramp current injections with continuous illumination of rising power densities during ramp; (**G**) Normalized light-induced delay of first spike on the ramp relative to the dark; 0 – no delay, 1 – no action potential during ramp; points represent individual recordings and lines averaged logistic fit; n=6, n=7; EC50 values mean±SD.

When we injected hyperpolarizing current steps to have more negative membrane potentials, we found that illumination caused depolarization instead of further hyperpolarization for both HcKCR1 and WiChR (Fig. 4D), explained by their residual Na^+^-conductance. However, reversion from depolarization to hyperpolarization occurred at a significantly more negative membrane potential of - 92±2 mV for WiChR than for HcKCR1 (−80±4 mV) (Fig. 4E, right) confirming the higher K^+^ -of WiChR.

Lastly, we used a somatic current ramp injection and increased illumination to compare the operational power densities at 1030 nm needed for effective inhibition with HcKCR1 and WiChR. (Fig. 4C) For both KCRs increasing power densities delayed the first action potential on the ramp compared to darkness, but half-maximal inhibition was observed at six times lower power densities for WiChR than for HcKCR1 (Fig. 4F,G, right). For HcKCR1 complete suppression of spiking on the ramp was achieved at 43 μW/μm^2^ while for WiChR, this was reached for all patched neurons already at 4.4 μW/μm^2^.

## Discussion

Potassium-selective channelrhodopsins are a new functional group of light-gated ion channels. In this study, we investigate their molecular mechanism and identify the amino acid motif that is required for their high potassium selectivity. Exploiting that knowledge, we uncovered two new potassium-conducting ChRs with further improved K^+^-selectivity, and in the case of WiChR with excellent inhibitory performance in neurons.

We show that potassium selectivity of KCRs relies on an unusual pair of aromatic residues in helix 3 and 7 that occludes the extracellular pore in the dark and constitutes the K^+^-selectivity filter in the open channel (W102 and Y222 in HcKCR1 highlighted in Fig 5A and Fig. S10), which is unique among rhodopsins. Additionally, six further residues are required for efficient K^+^-conductance. Among them, three pore-lining carboxylates (D87, D105 and D116 in HcKCR1) found at similar positions in ChRmine (*20*) are expected to be generally involved in channel gating, similarly as earlier discussed for the related GtCCR2 (*22*). Finally, three side chains of putative secondary importance (S73, F88 and L99 in HcKCR1) ensure conformational integrity of the pore and orientation and hydrophobicity of the K^+^-selectivity filter.

**Fig. 5:**
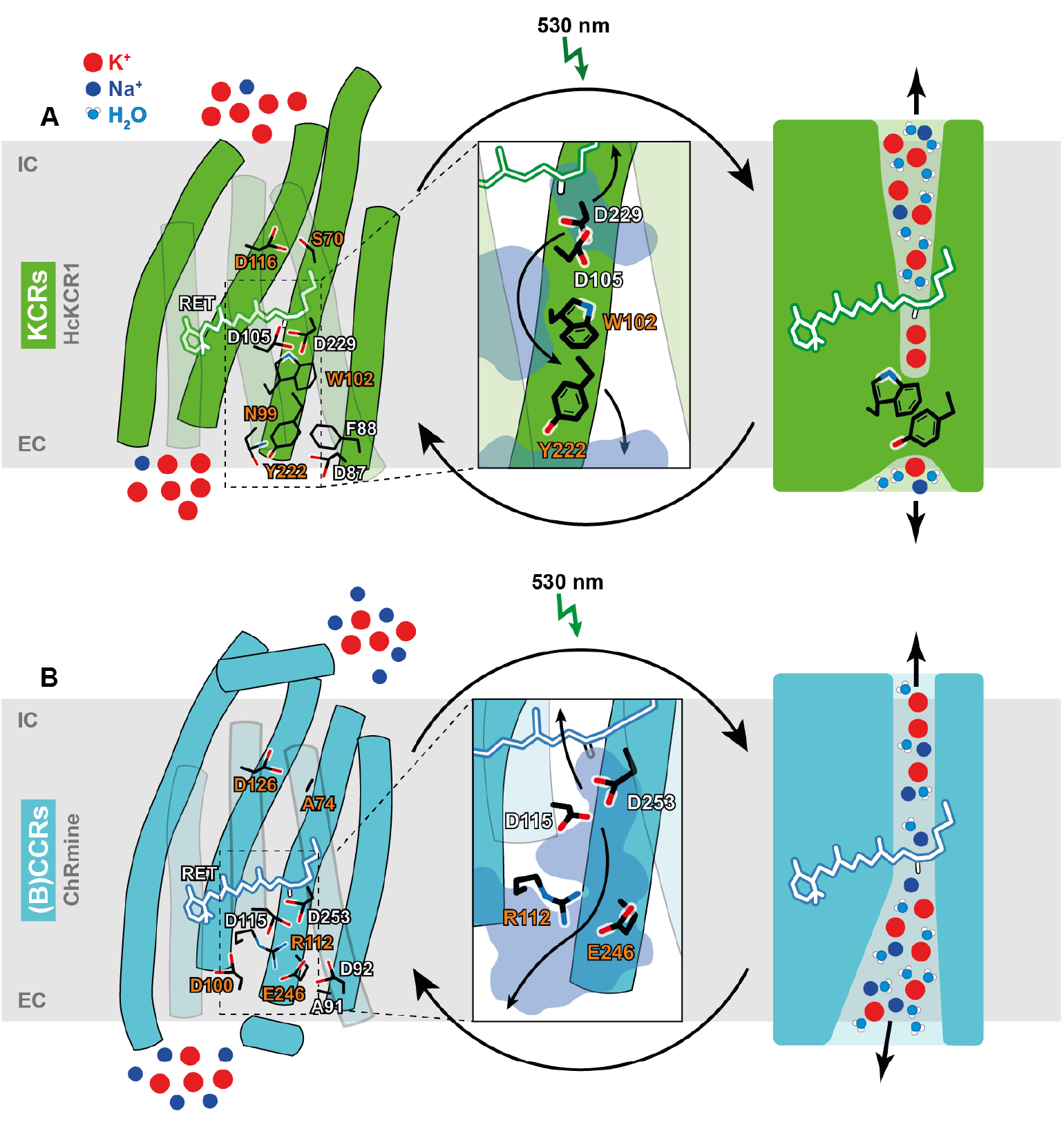
K^+^-selectivity filter of KCRs compared to (B)CCRs. (**A**)Cartoon model of the proposed HcKCR1 dark state and a simplified representation of its conducting state with important amino acid side chains highlighted. The central inlet shows the outer pore. Residues in light red are involved in K^+^-selectivity. (**B**)Cartoon model of the ChRmine dark state and a simplified representation of its conducting state. The central inlet shows the outer pore. Highlighted amino acids are homologous to residues in **A**. The outer pore in HcKCR1 is smaller and more isolated from the bulk than in ChRmine. Here, a hydrophilic and water-filled cavity between helices 3, 4, 6 and 7 is delimited from the central gate by W102 and Y106 and from the bulk water by W210, Y222 and N99. Geometrical RMSD calculations of aromatic residues revealed high flexibility of F88 (2.00 Å), W102 (2.12 Å), Y106 (1.85 Å) and W210 (2.27 Å) side chains while Y222 was more stable (1.13 Å) due to predicted hydrogen bonds to N99 and Q218. (Fig. S10B) This flexibility would allow the formation of a continuous cation translocation pathway during the conducting state.

Our findings stand in contrast to the highly conserved mechanism of ion conduction in conventional highly symmetric tetrameric K^+^-channels, like KcsA or Kv, where K^+^-ions are conducted along the tetramer interface tightly coordinated by backbone carbonyl oxygens of the selectivity filter (*23*). However, for KCRs, our MD simulation and mutational analysis predict a trimeric assembly with an inter-subunit cavity most likely filled by lipids similar to ChRmine (*19*) and an ion permeation pathway going through the individual KCR protomers themselves. We expect that here, the passage of potassium ions might require dehydration in order to pass the hydrophobic and aromatic side chains of the K^+^-selectivity where Na^+^ with higher dehydration energy and more localized charge is blocked. In the related ChRmine, the wider and presumably water-filled access funnel could allow permeation of partly hydrated K^+^ and Na^+^ (Fig. 5). A similar non-canonical mechanism for K^+^-selectivity was recently described for the lysosomal K^+^-channel TMEM175, that also features a hydrophobic restriction in the center of the pore as required for K^+^-conductance, with important differences in K^+^-selectivity for different channel subtypes ranging from *P*_K+_/*P*_Na+_ of 2 in bacteria to *P*_K+_/*P*_Na+_ of 35 in humans (*24*). Accordingly, *P*_K+_/*P*_Na+_ also varied significantly for different KCRs despite an overall conserved signature motif and we observe diverse conformations of the key residues along the pore in the different channels (Fig. S10).

Now, with the ability to control K^+^ flux in neuronal optogenetic experiments, KCRs will enable inhibition of excitable cells under yet inaccessible conditions and extend the possibilities beyond previously used ACRs. While effective neuronal inhibition is possible with HcKCR1, the newly identified WiChR, will outperform HcKCRs in several applications based on three superior properties. First the K^+^-selectivity over Na^+^ is more than three times higher for dark-adapted WiChR and even increases further over light-adapted HcKCR1 due to the low inactivation of WiChR. The increased K^+^-selectivity translates to neuronal application in our experiments and will allow effective inhibition of neurons over a broader range of resting potentials and less dependent on the Na^+^-gradient. Second, the high operational light sensitivity of WiChR allowed inhibition at an order of magnitude lower light doses compared to HcKCR1. On the one hand this will significantly facilitate light delivery for *in-vivo* 2P experiments and increase the accessible depth of neurons to inhibit, while on the other hand it allows the distribution of available laser power over a larger number of target cells and simultaneous inhibition of many cells. Third, the longer open state lifetime of WiChR is fundamental to the pulsed illumination protocol that we extensively investigated here and recommend for *in-vivo* application. Continuous illumination, as necessary for HcKCRs, will cause significant temperature rise in the tissue, especially considering the extended time frames and the number of cells to inhibit that can be needed for behavior experiments. On the contrary, the pulsed illumination can be applied indefinitely and for many target cells without any heat accumulation in the tissue presenting a tremendous advantage for *in-vivo* application. Lastly, we demonstrated a high activation of WiChR with targeted 2P holographic stimulation adding deeper tissue penetration, lateral single cell resolution and increased axial resolution over widefield 1P illumination, which likewise present considerable advantages for application.

Based on our results, we recommend WiChR as the tool of choice for targeted holographic 2P inhibition and simultaneous silencing of tens to hundreds of cells with low light intensities, short illumination times, in depth and with minimal photodamage.

## Materials and Methods

### Search for ChR genes

ChR genes from the stramenopile CCR subfamily: CbuChR1 (NCBI Genbank KAA0157615.1, NCBI genome assembly GCA_008330645.1) from *Cafeteria burkhardae* BVI (formerly, *Cafeteria roenbergensis* BVI), P1ChR1 (MMETSP1104_DN12643_c6_g1_i1, MMETSP assembly MMETSP1104) from Placidida sp. Caron Lab Isolate (formerly, *“Cafeteria”* sp.) and partial sequences of B1ChR1 and B1ChR2 from *Bilabrum* sp. (found as contamination in various algal transcriptome assemblies, in particular *Chroodactylon ornatum*, see Supplementary Text) were previously reported in (*4*). ChR genes of *Wobblia lunata* NIES-1015 (WlChR1-4) were obtained by assembling the raw data available from NCBI SRA (SRA run DRR049555, (*25*)) with Trinity v. 2.13.2 (*26*). Search for ChR genes in these and other assemblies was performed by querying protein sequences using microbial rhodopsin HMM profiles (https://github.com/BejaLab/RhodopsinProfiles) with hmmsearch from HMMER v. 3.3.2 (*27*). ChR sequences were obtained by searching against a curated database of microbial rhodopsins using blastp from blast+ v. 2.11.0+ (*28*). The complete list of searched stramenopile assemblies is provided in Suppl. Data S1. De novo assemblies when necessary were obtained with Trinity for transcriptomes or spades v. 3.14.1 (*29*) for genomes and single amplified genomes (SAGs). Genes were predicted with TransDecoder v. 5.5.0 for transcriptome assemblies and GeneMarkS v. 4.32 for genome and SAG assemblies. Additional sequences related to stramenopile CCRs were obtained from JGI freshwater and marine metatranscriptomes using blast with stramenopile CCRs as queries. Only publicly available complete sequences from JGI were used in the analyses.

A curated catalog of ChRs including sequences reported here is maintained at (*30*).

### Phylogenetic analysis

Phylogeny of the stramenopile CCRs, related clades and a reference set of ChRs from other families was obtained as follows. The sequences were aligned using mafft v. 7.475 (*31*) in --localpair mode, trimmed with trimAl v. 1.4.1 (*32*) (with -gt 0.9) and the phylogeny was reconstructed with iqtree2 v. 2.1.2 (*33*) with 1000 ultrafast bootstrap replicates (*34*). The tree was outgroup-rooted.

### Structure-based alignment

Representative CCRs were aligned with 3D-coffee from t_coffee v.13.45.0.4846264 (methods sap_pair, mustang_pair, t_coffee_msa and probcons_msa) (*35*). Transmembrane regions in ChRmine (PDB: 7W9W) were predicted with the PPM web server (*36*).

### Molecular Biology and Cell Culture

Coding sequences of HcKCR1 (MZ826862), HcKCR2 (MZ826861), B1ChR2 and WlChR1 (synthesized by GeneScript) and ChRmine (gently provided by K. Deisseroth) were fused to a Kir2.1 membrane targeting sequence, a mScarlet, mCerulean or eYFP fluorophore and an ER-Export signal and cloned either into a pCDNA3.1 vector behind a CMV-promotor for basic characterization in ND7/23 cells or into a pAAV backbone behind a human Synapsin promotor for virus production and expression in neurons. Site-directed mutagenesis was performed using a QuickChange Site-Directed Mutagenesis Kit (Agilent Technologies, Santa Clara, CA), according to the manufacturer’s instructions. Expression in ND7/23 cells was performed as previously described (*37*). ND7/23 cells were cultured at 5% CO_2_ and 37 °C in Dulbecco’s minimal essential medium supplemented with 5% fetal bovine serum, 1 μM all-*trans* retinal and 100 μg/ml penicillin/streptomycin (Biochrom, Berlin, Germany). Cells were seeded on poly-lysin coated coverslips at a concentration of 0.75 × 10^5^ cells/ml and transiently transfected using the FuGENE^®^ HD Transfection Reagent (Promega, Madison, WI) 28–48 h before measurement.

### Whole-Cell patch clamp in ND7/23 cells

Patch pipettes were prepared from borosilicate glass capillaries (G150F-3; Warner Instruments, Hamden, CT) using a P-1000 micropipette puller (Sutter Instruments, Novato, CA) and were subsequently fire-polished. Pipette resistance was 1.5 to 2.5 MΩ. A 140 mM NaCl agar bridge served as reference (bath) electrode. In whole-cell recordings, membrane resistance was typically > 1 GΩ, while access resistance was below 10 MΩ. Pipette capacity, series resistance, and cell capacity compensation were applied. All experiments were carried out at 23 °C. Signals were amplified and filtered at 2 kHz, digitized at 10 kHz (DigiData1400) and acquired using Clampex 10.4 Software (all from Molecular Devices, Sunnyvale, CA). Continuous monochromatic light was generated using a Polychrome V light source (TILL Photonics, Planegg, Germany) coupled into the Axiovert 100 microscope (Carl Zeiss) and delivered to the sample using a 90/10 beamsplitter (Chroma, Bellows Falls, VT). Light exposure was controlled by a VS25 and VCM-D1 shutter systems (Vincent Associates, Rochester, NY). Light intensities were adjusted either manually by different neutral density filters or by a motorized Filter-wheel for equal photon densities in action spectra. Light intensities were measured in the sample plane with a calibrated P9710 optometer (Gigahertz Optik, Türkenfeld, Germany) with 3.7 mW/mm^2^ at 470 nm and 2.7 mW/mm^2^ at 530 nm for standard measurements. The illuminated field of the W Plan-Apochromat 40x/1.0 DIC objective was 0.066 mm^2^ (Carl Zeiss, Jena, Germany).

Extracellular buffer exchange was performed manually by adding at least 5 mL of the respective buffer to the recording chamber (500 μL chamber volume) while a Ringer Bath Handler MPCU (Lorenz Messgerätebau, Katlenburg-Lindau, Germany) maintained a constant bath level. Standard bath solutions contained 110 mM NaCl, 1 mM KCl, 2 mM CaCl2, 2 mM MgCl2, and 10 mM HEPES at pH_e_ 7.2 (with glucose added up to 310 mOsm). Standard pipette solutions contained 110 mM Potassium-D-gluconate, 1 mM NaCl,, 2 mM CaCl_2_, 2 mM MgCl_2_, 10 mM EGTA, and 10 mM HEPES at pH_i_ 7.2 (glucose was added up to 290 mOsm). For ion selectivity measurements, extracellular NaCl was replaced by 110 mM KCl or 110 mM NMDGCl.

I(E)-curves were measured from −80 mV to +40 mV in 20 mV steps with liquid junction potential (LJP) corrected holding voltages and 500 ms illumination. Action spectra were recorded with 110 mM Na^+^_e_, 10 ms illumination and reduced light intensities at either 0 mV for K^+^-selective channels and mutants or −60 mV for HcKCR1 mutants that lost K^+^-selectivity. Light titrations of KCR channels were recorded at 0 mV with 500 ms illumination.

ND7/23 measurements were analyzed using the Clampfit 10.7 software (Molecular Devices, Sunnyvale, CA), Microsoft Excel and Origin 2020^®^ (OriginLab, Northampton, MA). Photocurrent traces were baseline corrected, filtered, and reduced in size for display purposes. Photocurrents were normalized to peak photocurrents at - 40 mV and 110 mM K^+^e and action spectra were fitted using a parametric Weibull function:

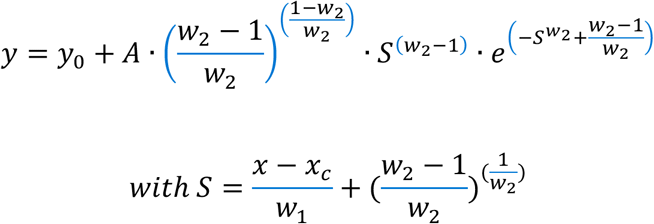

and the estimated parameters A, yo, wi, w2, xc.

### Confocal microscopy in ND7/23 cells

Confocal images were acquired using a FluoView1000 (Olympus, Tokyo, Japan) confocal laser scanning microscope, with a UPlanSApo 60x NA 1.2 water-immersion objective (Olympus, Tokyo, Japan). mScarlet was excited with a 559 nm DPSS laser (2% transmissivity), mCerulean with a 440 nm diode laser (10% transmissivity), and eYFP was excited with an argon laser (3% transmissivity).

Both the membrane fluorescent density (brightness/area) and the fluorescent density within the cell were calculated from equatorial slices after background subtraction. The membrane was defined as the outermost layer of the cell, and the targeting was then evaluated as the fluorescence membrane density in the membrane divided by the intracellular fluorescence density. This calculation yields a value below one if the fluorescence is higher inside the cell, and above one if the membrane is brighter.

### Preparation and electroporation of organotypic hippocampal slices

All animal procedures were performed in accordance with French law and guidelines of Directive 2010/63/EU and institutional guidelines on care and use of laboratory animals. Organotypic hippocampal slices were prepared from P5/P8 C56BL/6J mouse pups as described in Gee et al. – 2017 (*38*). In brief, dissection of hippocampi was carried out in ice cold dissection solution composed of: 248 mM sucrose, 26 mM NaHCO_3_, 10 mM glucose, 4 mM KCl, 5 mM MgCl_2_, 1 mM CaCl_2_, 2 mM kynurenic acid and 0.001% phenol red saturated with 95% O_2_/5% CO_2_. 400 μm thick transverse sections of hippocampi were sliced with a McIlwain Tissue Chopper using double edge stainless steel razor blades. Morphologically nice slices were carefully transferred using a plastic transfer pipette onto small pieces of PTFE membrane (Millipore FHLP04700) on membrane inserts (Millicell PICM0RG50) in pre-warmed culture medium. The slices were cultured at 37°C and 5% CO_2_ in culture medium consisting of 80% MEM and 20% Heat-inactivated horse serum supplemented with 1 mM L-glutamine, 0.01 mg/ml Insulin, 0.00125% Ascorbic acid D-glucose, 14.5 mM NaCl, 2 mM MgSO_4_, and 1.44 mM CaCl_2_; no antibiotics were added to the culture medium. Three to four slices were cultured on one insert and every 3 days the medium was partially exchanged with pre-warmed fresh medium.

Bulk electroporations were performed with endotoxin-free plasmid preparations (>0.9 μg/μl) using the Y-unit of the 4D-Nucleofector^®^ (Lonza Bioscience), both HcKCR1 and WiChR were subcloned into a pAAV-backbone and expressed as fusion proteins tagged with a fluorophore under a human syn1 promoter. At DIV3-4, the hippocampal slice was placed in a well of a 24-well culture plate (Nunc) and 10 μg plasmid DNA in a volume of 25-30 μL of AD1 Solution (Lonza Bioscience) was put as a drop on the slice. After an incubation period of 20 minutes at 37 °C, pre-warmed AD1 solution was added to a final volume of 360 μL. The 24-well Dipping Electrode Array (Lonza Bioscience) was carefully placed onto the 24-well culture plate avoiding air bubbles, which would disrupt the electroporation, immediately followed by the electroporation using program CM-158 of the 4D-Nucleofector® Y Unit. The slices were used for electrophysiology recordings 6-10 days after electroporation.

### Holographic microscope for two-photon excitation

The system, capable of holographic light shaping (*2*), is schematically represented in Fig. S11A. It was custom-built around a commercial upright microscope (Zeiss, Axio Examiner.Z1). An efficient holographic stimulation was based on the use of a low repetition rate amplified fiber laser, providing ~300 fs pulses at 500 kHz rate with a 1030 nm wavelength (Satsuma, Amplitude Lasers, total output power =10 W). The laser beam, after a first beam expander (f1 = 50 mm; f2 = 200 mm), was directed on a spatial Light Modulator (SLM) (LCOS-SLM X13138-07, Hamamatsu Photonics, 1280 × 1024 pixels, 12.5 μm pixel size). Then, a couple of 4f system (f1= 750 mm and f2 = 250 mm; f3 = 50 mm and f4 = 250 mm), projected the SLM plane to the pupil of a 20x 1.0 NA objective (Zeiss, W Plan-Apochromat 20x/1.0 DIC). The SLM was controlled by custom-made C++ based software (*39*) that uses a variation of the Gerchberg and Saxton algorithm (*40*) in order to calculate a phase-hologram to modulate the laser beam and to generate an arbitrary illumination shape at the sample plane. For single neuron excitation, a 12 μm-diameter circular spot was generated. A physical blocker was placed between the two lenses of the first 4-f system to avoid the zero-order of diffraction reaching the sample. For alignment and calibration, holographic patterns were projected on a thin spin-coated rhodamine fluorescent layer and the induced fluorescence was visualized on a CMOS camera (Hamamatsu ORCA-5G). To reconstruct a full 3D profile of the illumination spot (as reported in Fig. S11), a second inverted microscope was built in transmission geometry and different axial sections of the spot were imaged on a second bottom camera while translating axially the upper objective (as described in (*40*)). A fast shuttering and power control of illumination pulses were achieved using a built-in acousto-optic modulator that was controlled by the Digitizer (DigiData 1440, Molecular Devices) to ensure synchronization with the electrophysiology recordings. The microscope was additionally capable of widefield fluorescence imaging based on a multi-led illumination system (pE-4000, CoolLed) and a DIC transmitted imaging.

For whole-cell patch-clamp recordings the microscope is equipped with a micromanipulator (Junior XR, Luigs and Neuman); signals were amplified and digitized using an AxoPatch 700B (Molecular Devices) and a DigiData 1440A (Molecular Devices) while data was acquired using Clampex (Molecular Devices).

### 2-Photon electrophysiology recordings in organotypic slices

Recordings were performed 6-9 days after electroporation with transfected cells identified by their fluorescence. Patch pipettes were pulled from fire-polished capillaries with filament (OD 1.5 mm/ ID 0.86 mm, World Precision Instruments) using a micropipette puller (P1000, Sutter Instruments). Pipettes with a resistance in a range from 3-6 MΩ were filled with a K-gluconate based intracellular solution (in mM): 135 K-Gluc, 10 HEPES, 10 Na-Phosphocreatin, 4 KCl, 4 Mg-ATP, 0.3 Na2-GTP; 290 mOsm. Extracellular solution was composed of (in mM): 125 NaCl, 26 NaHCO_3_, 1.25 NaH_2_PO_4_, 25 D-Glucose, 2.5 KCl, 1.5 CaCl_2_, 1 MgCl_2_, 0.5 ascorbic acid (310 mOsm), continuously bubbled with carbogen (pH 7.4) and perfused at a rate of 1.5-2.5 ml/min (room temperature). The liquid junction potential was calculated to be – 17.7 mV and corrected post acquisition for all recordings; the Nernst potential for potassium is −103 mV while that for sodium is +48 mV in these conditions.

After establishing the whole-cell configuration in voltage clamp mode, quality of the patch was checked and access resistance was continuously monitored during and after the recording (voltage clamp); if access resistance rose above 45 MΩ recordings were discarded. After switching to current clamp mode bridge balance compensation was performed with the internal circuits of the amplifier and neuronal parameters like resting potential, input resistance and rheobase were determined (Fig. S12B). During the recording constant holding currents (between −50 pA and +50 pA) were injected (if necessary) to keep the neurons at a comparable membrane potential between −75 mV and −80 mV.

For the 2-photon stimulation, a circular holographic spot of ≈12 μm diameter (Fig. S12B) was generated aiming at the cell body of the patched neuron. Stimulation light powers were measured every recording day out of the microscope objective (PM160 power meter, Thorlabs), fitted within the range of interest and used to calculate the exact photostimulation intensity; a typical average spot size of 126 μm^2^ was determined from multiple images of the spot fluorescence on a thin rhodamine layer and used to calculate the power densities reported in the respective figures.

Square current injections of 1 s duration were increased in 10 pA steps per sweep starting with hyperpolarizing injections, with a 5 s pause between sweeps. The duration of the current injection was divided into the first 250 ms (before illumination), 500 ms (during illumination) and 250 ms (after illumination) for analysis. Illumination was either continuous for the whole 500 ms or a sequence of 5 ms pulses at different frequencies (30-35 μW/μm^2^). The firing rates were determined during these intervals and normalized to the firing rate during the first 250 ms. The firing rates were extracted for each neuron at current injections 50 pA and 100 pA above the respective rheobase of the neuron; with the rheobase determined as the current injection were the first spike in the first or last 250 ms (no illumination) occurred. After extraction the normalized firing rates were averaged and plotted as mean ± SD. Control recordings were performed and analyzed in the same way, but without any light application in the middle 500 ms.

For recording the reversal from light-induced hyperpolarization to depolarization, square current injections were increased in 10 pA steps around the reversal and after 250 ms continuous illumination at saturating intensities (30-35 μW/μm^2^) was switched on. Light-induced ΔV_mem_ was extracted as the difference between V_mem_ before illumination and at 150 ms into the illumination; the value for the reversal was extracted by linear interpolation on the individual recordings and then averaged over all cells.

For the comparison of the effectiveness of inhibition somatic ramp current injections (800 ms) with continuous illumination of rising power densities were applied (5 s waiting between sweeps). The amplitudes of the ramp were constant for each individual neuron and chosen to give 10-20 action potentials in the dark respectively; between 75 pA to 500 pA. The timing of the action potential was extracted at the peak amplitude of the spike and the delay was calculated with respect to the first action potential on the ramp injection without illumination. Afterwards, the delay was normalized between no delay (0) and maximum delay (1, no spike on ramp). Resulting values were plotted against the power densities of the respective recording day and fitted with a five parameter logistic function; then values of the individual recordings were averaged.

Data was analyzed using Clampfit, custom Python scripts, Easy Electrophysiology and Origin 2021.

### Temperature Simulations

The temporal distribution of the temperature rise reported in Fig. S13 was calculated by solving the Fourier heat diffusion equation (*41*) considering the brain tissue as an infinite medium with isotropic and uniform thermal properties as described and experimentally verified before (*42*). For these simulations light scattering in the tissue was neglected.

The reported temperature rise was calculated for r = 0 representing the center of the first holographic spot in the middle of the field of view. Additional spots were randomly added to the first one to maintain a relatively uniform distribution within the 100 x 100 μm surface.

### 3D structure prediction and system preparation for MD simulations

The HcKCR1, WiChR and B1ChR2 3D models were generated using their full-length amino acid sequences with AlphaFold2 v.2.1.1 (*43*)) with the full structural dataset running on a dedicated group server. Based on the sequence similarity to ChRmine, a trimeric assembly was chosen. For ChRmine, the trimeric cryo-electron microscopy structure was used (PDB-ID 7SFK) (*19*). Each protomer of HcKCR1, WlChR1 and B1ChR2 was equipped with a retinal cofactor bound to K233 in HcKCR1, K251 in WlChR1 and K232 in B1ChR2. Internal hydration was predicted using Dowser++ (*44*). The central pores of the HcKCR1, WiChR, B1ChR2 and ChRmine trimer were closed using 1,2-dimyristoyl-sn-glycero-3-phosphocholine (DMPC) molecules manually placed in PyMOL v.2.5.0 (Schrödinger, LLC). Accordingly, three lipids were positioned for HcKCR1, WiChR, ChRmine and 6 for B1ChR2. All systems containing the protein, internal waters and central lipids were introduced into a homogeneous DMPC bilayer membrane, respectively, surrounded by water and neutralized using 150 mM KCl using CharmmGUI (*45*).

### Classical MD simulations in CHARMM at constant pH

MD simulations were carried out using CHARMM version c42b1 (*46*) and openMM version 7.0rc1 (*47*). The MD simulation was computed under NPT conditions using a 2 fs time step, a 303.15 K Langevin heat bath, the particle-mesh Ewald method for long-range electrostatics, and the CHARMM36 force field (*48*). Before and between the initial equilibration steps, the protonation of all titratable residues was adjusted based on periodical Karlsberg2^+^ predictions (*49, 50*). For further production runs, combined pKa-MD simulations were computed: during classical MD simulations, pKa values were calculated periodically for 11 snapshots of each 10 ns production run using Karlsberg2+MD (*50*) to subsequently adapt the protonation pattern of the system pH dependently (pH 8).

### pK_a_ calculations in Karlsberg2^+^

In Karlsberg2^+^, the holoprotein structure and ions within a 4 Å cutoff were explicitly included. The rest was substituted with continuum solvation and an implicit ion concentration of 100 mM. Single-structure pKa calculations during the equilibration were performed using APBS (*51*) in a conformational space of three pH-adapted conformations (PACs), while the pKa calculations based on MD simulations used the extracted snapshots and created only single PACs at pH 7. All PACs were generated using Karlsberg2^+^ in a self-consistent cycle including adjustment of protonation patterns of titratable amino acids and salt bridge opening according to either pH −10, 7, or 20.

### Statistical Analysis

Throughout the paper data is shown as mean ± standard deviation; additionally individual data points are displayed. Number of replicates and statistical analysis (if performed) are mentioned in the respective figure legends.

## Acknowledgments

We thank Yinth Andrea Bernal Sierra for helpful discussions and Aysha Mohammed Lafirdeen for the preparation of organotypic slices.

## Funding

Deutsche Forschungsgemeinschaft (DFG, German Research Foundation) - SPP1926 - 425994138 (PH);

Deutsche Forschungsgemeinschaft (DFG, German Research Foundation) under Germany’s Excellence Strategy – EXC-2049 – 390688087 (JV);

European Research Grant Stardust, No. 767092 (PH, EP);

French National Research Agency ANR Holoptogen AAPG2019 (VE, PH);

Israel Science Foundation grant 3592/19 and 3121/20 (OB);

Deutsche Forschungsgemeinschaft (DFG, German Research Foundation) 442616457 (CG);

PH is a Hertie Professor supported by the Hertie Foundation;

OB holds the Louis and Lyra Richmond Chair in Life Sciences

## Author contributions

Conceptualization: PH, JV, EP and CG

Bioinformatics: AR and OB

Electrophysiology: JV, EP and SA

Neuronal studies: CG, DT and VE

Simulations: EP and BCF

Confocal microscopy: ACS

Funding acquisition: PH, OB, VE and JV

Visualization: JV, EP, CG and AR

Writing: JV, CG, EP, AR and PH with contributions from all authors.

## Data and materials availability

Annotated transcript sequences of the stramenopile CCR genes from *Bilabrum* sp., Placidida Caron Lab Isolate and *Wobblia lunata* are provided in Suppl. Data File S3

## List of Supplementary materials

Supplemental Text

Supplemental Fig. S1 - S13

Supplemental Data File S1. Transcriptome and genome assemblies of Stramenopiles used for searching ChR genes. Excel file.

Supplemental Data File S2. Rhodopsin sequences used for phylogeny reconstruction and the resulting unrooted tree. Branch supports are ultra-fast bootstrap support values. Zip file containing fasta and newick files.

Supplemental Data File S3. Annotated transcript sequences of stramenopile CCR genes from *Bilabrum* sp., Placidida Caron Lab Isolate and *Wobblia lunata*. Zip file containing genbank flat files.

Supplemental Data File S4. Trimmed structure-based alignment of ChRmine, HcKCR1, HcKCR2, WlChR1 and B1ChR2. Clustal alignment file.

Supplemental Data File S5-S7. Equilibrated AlphaFold2 model of the HcKCR1, WiChR and B1ChR2 trimeric assemblies. PDB files.

