## Supplementary Materials for "WiChR, a highly potassium selective channelrhodopsin for low-light two-photon neuronal inhibition"

**This PDF file includes:**

Supplementary Text
Figs. S1 to S13
References (1 to 9)

**Other Supplementary Materials for this manuscript include the following:**

Data S1to S5

#### Supplementary Text

<u>The origin of B1ChR1 and B1ChR2</u>

Two partial genes from the stramenopile CCR clade, 1KP_LLXJ_2038500 and 1KP_LLXJ_2042068, were extracted by us previously from the transcriptome assemblies of the red alga *Chroodactylon ornatum* (1000 Plant transcriptomes [1KP] assembly LLXJ) and the green alga *Acrosiphonia* sp. SAG 127.80 (1KP assembly JIWJ)(*1*). Together with a third gene related to *Cafeteria* ACRs (1KP_LLXJ_2001886 and 1KP_LLXJ_2001887), nearly identical ChRs were found in transcriptome assemblies of two additional unrelated algae: *Halochlorococcum marinum* (ALZF) and *Picochlorum atomus* (JQFK) signifying that none of the four algae were the source of the ChR genes. To clarify the origin of the ChRs, 18S rRNA fragments were extracted from the four assemblies by searching the transcriptomes with blastn from blast+ using *Homo sapiens* 18S as the query. The matched sequences longer than 250 nt were searched against the nr database and the only shared 18S gene shared by all four assemblies matched *Bilabrum latius* HFCC35 SSU rRNA (MN315515.1) with identity values ranging from 99-100% (Fig. S7B). Search against a collection of algal and protist protein database with ublast from usearch v. 11.0.667 (https://drive5.com/usearch/) with an e-value threshold of 1e-10 and recording best-matching taxa for each protein in the four assemblies indeed revealed a sizable fraction of transcripts of stramenopile origin (Fig. S7C). Thus, we attributed the source of the common contamination in the four algal cultures to a non-clonal *Bilabrum sp.* (Anoecida) and the ChRs genes were labeled B1ChR1 (1KP_LLXJ_2038500), B1ChR2 (based on 1KP_LLXJ_2042068, see below) and B1ChR3 (1KP_LLXJ_2001886). Further support to the anoecid origin of B1ChR1-3 is provided by the close relationships of B1ChR1 to CbuChR1 (members of the stramenopile CCR clade) and B1ChR3 to CarACRs (collectively: anoecid ACRs) from the other anoecid *Cafeteria burkhardae* (Fig. S6A). The two species are the only currently known stramenopiles with genes from both of these ChR families.

The truncated 5’ end of the B1ChR1 transcript was extended by iterative read mapping with bowtie2 v. 2.3.5.1 (*2*) in --sensitive-local mode using the raw data in SRA run ERR2041128. The original transcript 1KP_LLXJ_2042068 and some of other B1ChR2 variants appeared to contain premature stop codons. Raw read mapping from the four source datasets revealed the presence of stop-free variants in SRA runs ERR2041020 and ERR3487419. Thus, the raw reads in ERR2041020 were mapped to the transcript 1KP_LLXJ_2042068 using bowtie2, duplicate reads were removed with MarkDuplicates from picard v. 2.26.5 (http://broadinstitute.github.io/picard/) and phased with samtools phase v. 1.11 (*3*). The haplotype with no premature stop codons was chosen and the majority-rule consensus sequence was taken to represent the gene.

<u>Distribution of ChRs among stramenopiles</u>

Systematic search for ChR genes in various algal and protist groups (see Suppl. Dataset S1) revealed that the KCRs reported from Hyphochytrium catenoides (HcKCR1, HcKCR2 and HcChR3) are part of a clade (Figs. S6A and S7A). The other stramenopiles with members of this ChR group are marine heterotrophic flagellates from the clade of Opalozoa: *Wobblia lunata* and *Placidida sp.* Caron Lab Isolate (Placidida) and *Cafeteria burkhardae* and *Bilabrum sp.* (Anoecida) (Figs. S6A and S7A). Additional related sequences were obtained from metatranscriptomic datasets. Given the fact that the successfully characterized members of the clade are cation (potassium) channels, we designate the family as stramenopile CCRs. Closely related to stramenopile CCRs are cryptophyte CCRs and a novel clade of uncharacterized ChRs from colpodellid alveolates. A more basal position is occupied by dinoflagellate channels whose selectivity is not yet known (*4*) and a yet another clade of ChRs from colpodellids. The opposite branch of the ChR tree is occupied by all of the known ACR families including three previously characterized families from Stramenopiles and a singleton ChR from MAST-22 and the monophylum uniting prasinophyte ACRs and green algal CCRs (Fig. 3B, S6A). It is curious to note that among Stramenopiles only the anoecids appear to have ChRs from both the CCR clade and an ACR clade while the other groups possess only one ChR type each. Given the presence of related ChR families in the other phylum of the SAR supergroup, the alveolates (dinoflagellates and colpodellids), it is tempting to hypothesize that the stramenopile CCRs were present in the last common ancestor of the phylum and were lost many times, most notably in the lineage of photosynthetic stramenopiles (Ochrophyta), although lack of strong phylogenetic signal precludes robust testing of this hypothesis. The obtained clustering pattern of the stramenopile CCRs might indicate that the common ancestor of Anoecida and Placidida had two CCR genes and that *H. catenoides* acquired its ChRs from Opalozoan, although this remains to be replicated in the future on a bigger sample of species and proteins. As for other ChRs in general, physiological role of the different ChRs in stramenopiles is hypothesized to be in phototactic reactions in flagellated life stages (swarmers in the case of labyrinthulids (*5*) and H. catenoides).

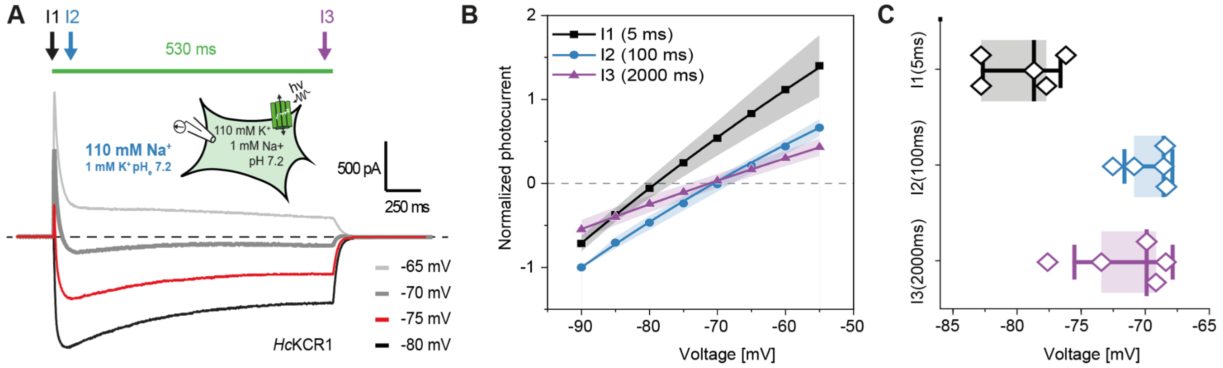


**Fig. S1. $K^+$-selectivity changes of HcKCR1 during continuous illumination. (A)** Photocurrent traces recorded at 4 selected voltages. **(B)** I(E) relation for the three time points $I_1$, $I_2$, and $I_3$ either 5 ms, 100 ms or 2000 ms after light on and **(C)** the three determined $E_{rev}$ values. Note that the selectivity adapts during the first 100 ms, whereas the current amplitude continues to decline for another 2 seconds until $I_3$. Data show Mean ± SD with n = 5.

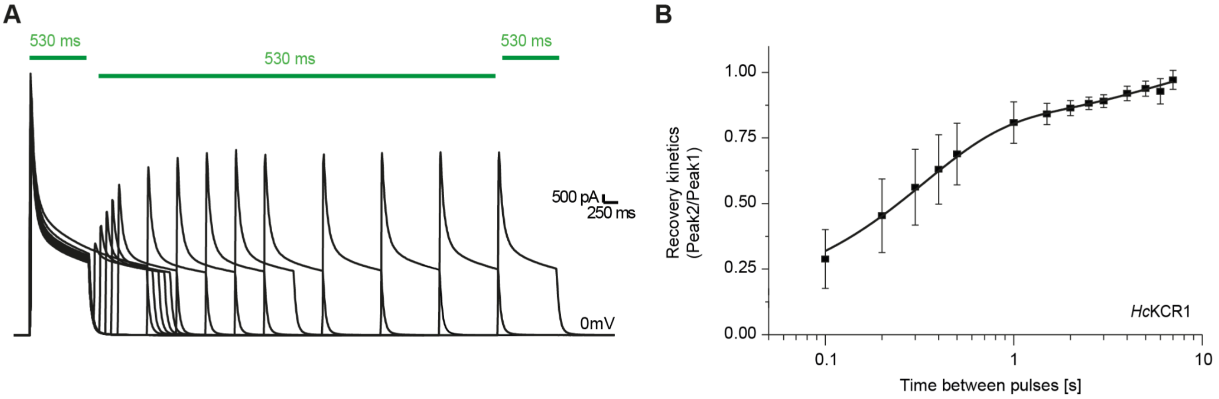


**Fig. S2. HcKCR1 photocurrent adaptation and recovery in darkness. (A)** A set of 14 double pulse experiments with increasing delay between pulses. **(B)** Recovery kinetics as the peak ratio of peak2 over peak1 plotted against the second pulse delay (Mean ± SD, n = 5). Photocurrents were measured at 0 mV with 110 mM $Na^+_e$ and 110 mM $K^+_i$.

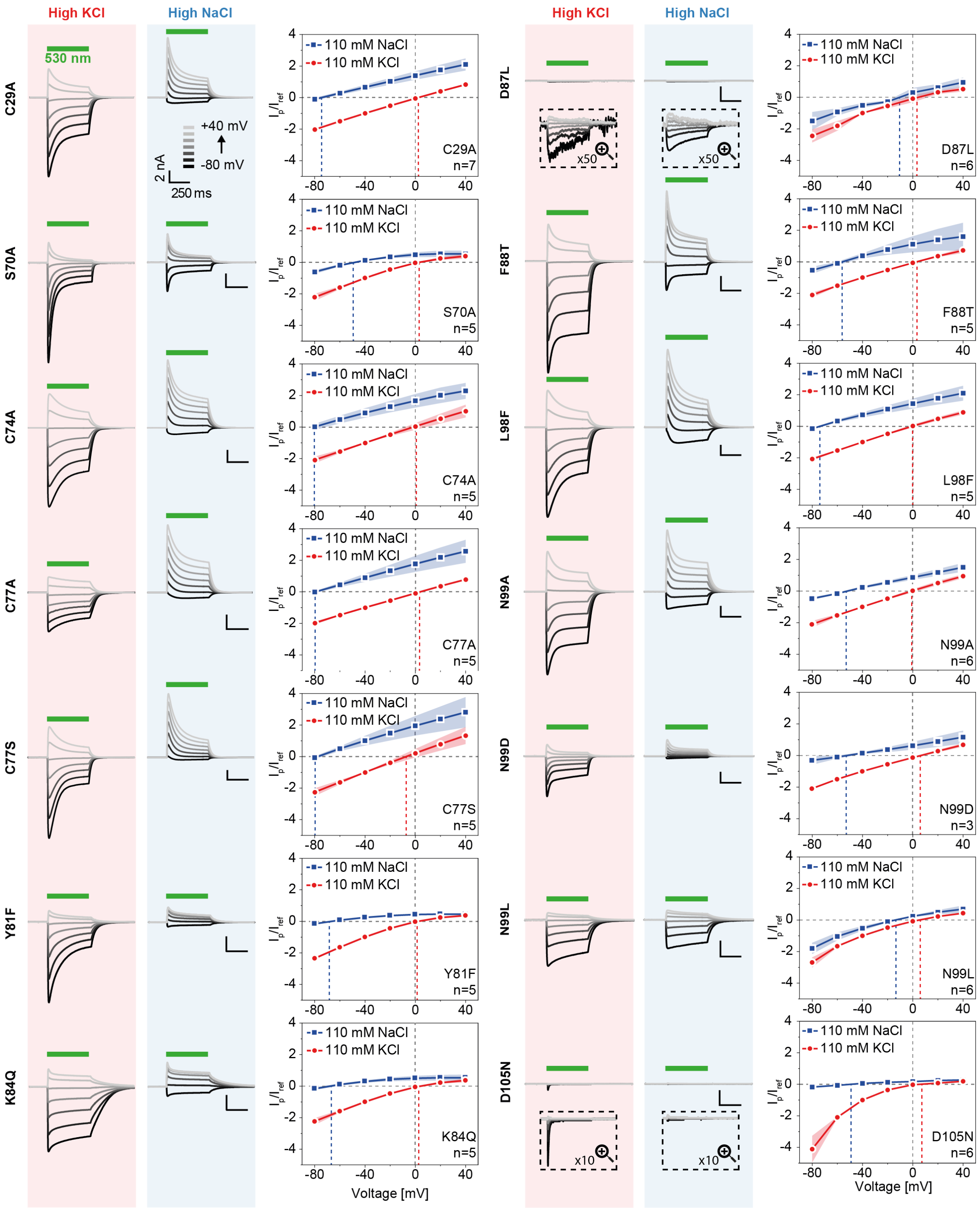


**Fig. S3.** Representative photocurrent traces at either high $K^+_e$ (red) or $Na^+_e$ (blue) and $I(E)$ relations of HcKCR1 pore mutants with 530 nm illumination. Boxes describe magnified signals with the given magnification factor. Most mutations have no to little influence on the $K^+$ selectivity except for D87L and N99L. All data show mean ± SD.

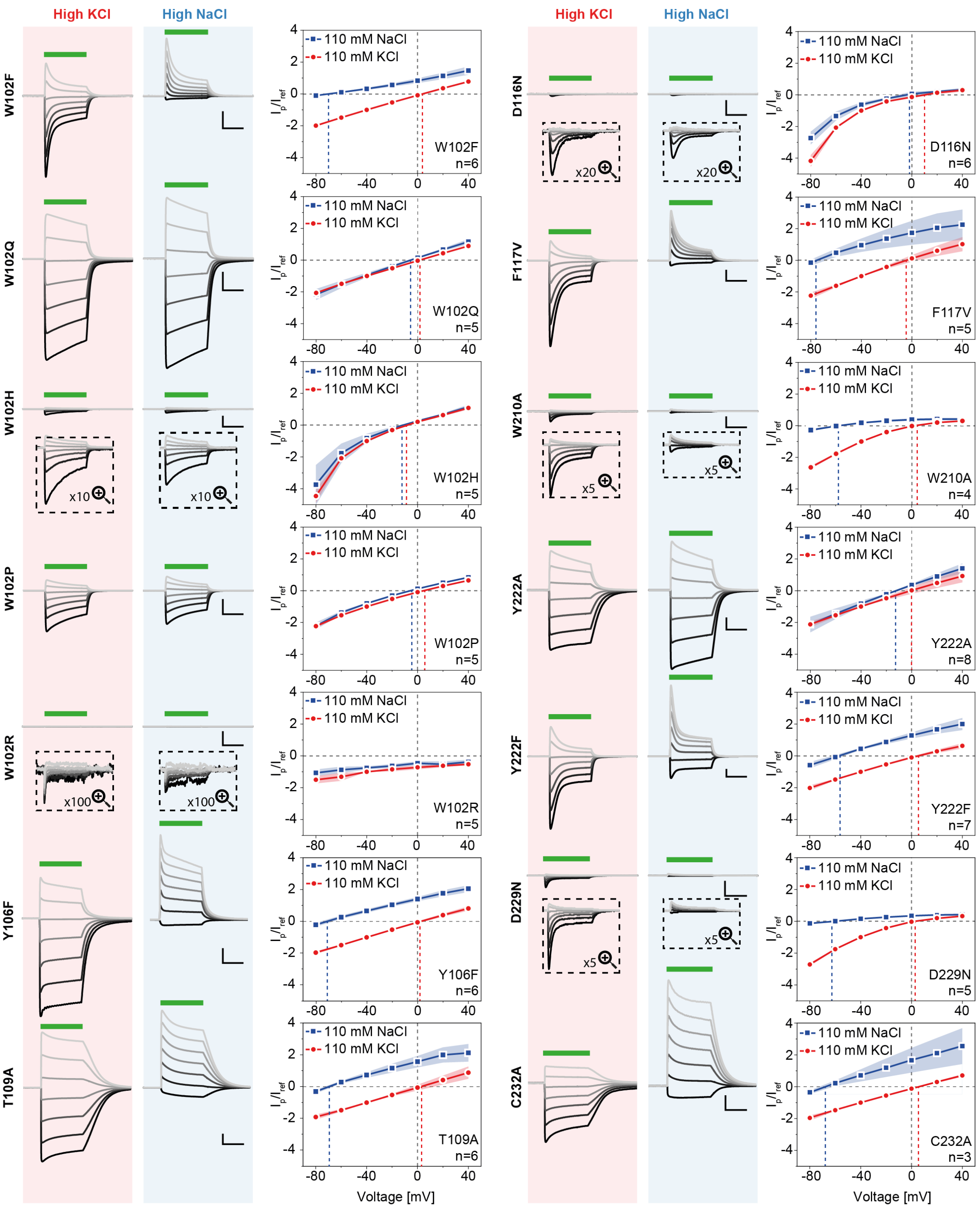


**Fig. S4.** Photocurrent traces at either high $K^+_e$ (red) or $Na^+_e$ (blue) and *I*(*E*) relations of HcKCR1 pore mutants illuminated with 530 nm. (continued). W102 mutations to Q, H and P as well as D116N and Y222A show drastic $K^+$ selectivity loss. All data show mean ± SD.

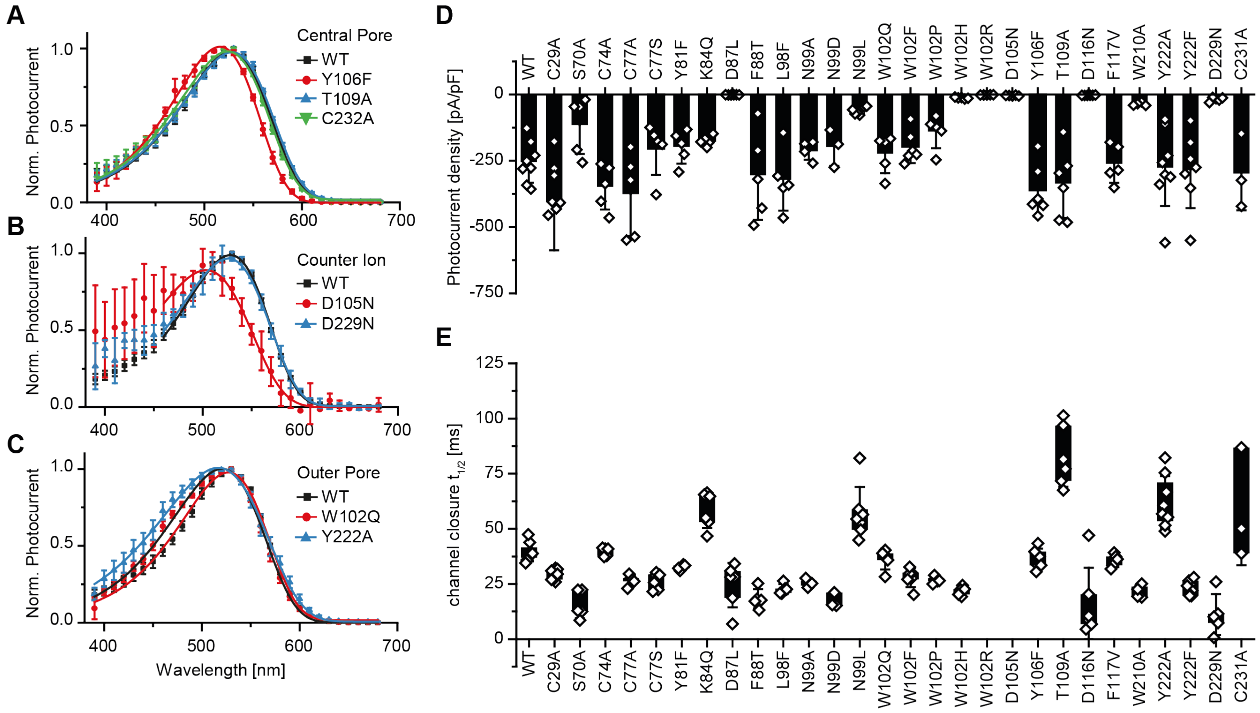


**Fig. S5. Spectral and electrophysiological properties of HcKCR1 and pore mutants. (A)** Action spectra of normalized peak photocurrents following 10 ms light pulses of different color but equal photon density in for HcKCR1-WT and central pore mutants (n = 6/6/7/5 for WT/Y106F/T109A/C232A at 0 mV) **(B)** for counter ion mutants (n = 6/6/5 for WT/D105N/D229N at 0 mV) and **(C)** for outer pore mutants (n = 6/5/9 for WT/W102Q/Y222A at -40 mV). (**D**) Peak-photocurrent densities at -40 mV holding voltage and **(E)** channel closure half time at -40 mV holding voltage. Photocurrents in (A) were recorded with high extracellular $Na^{+}_{e}$ and in (B) and (C) with high extracellular $K^{+}_{e}$. All data show mean ± SD.

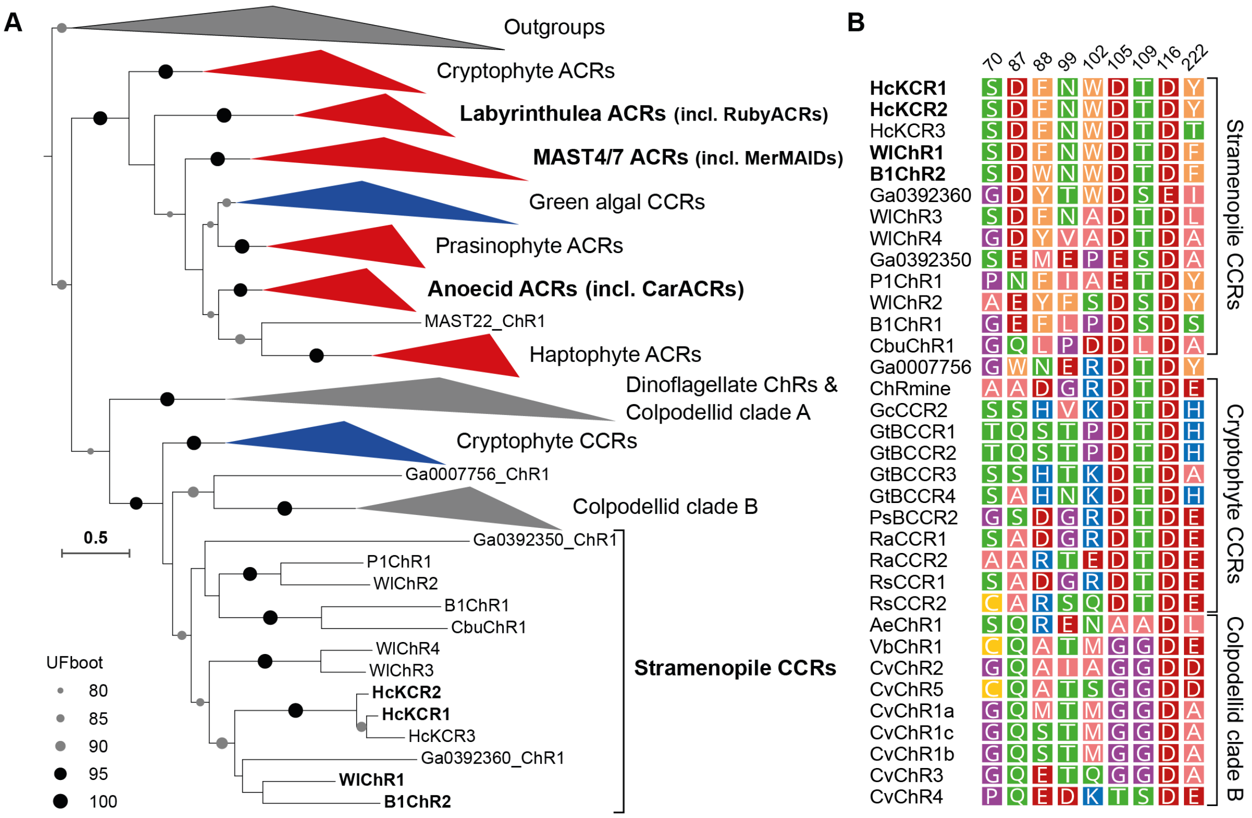


**Fig. S6. Phylogenetic analysis and diversity of the stramenopile CCRs and related ChRs.** **(A)** Phylogeny of channelrhodopsins. ChR subfamilies are colored by the selectivity of the characterized members: blue for cation channels and red for anion channels. Highlighted in bold are tested stramenopile CCRs expressed before (HcKCR1, HcKCR2, WlChR1 and B1ChR2). Species prefixes are as follows: B1 - *Bilabrum* sp., Cbu - *Cafeteria burkhardae*, Hc - *Hyphochytrium catenoides,* P1 - Placidida sp. Caron Lab Isolate, Wl - *Wobblia lunata*. The sequences and the tree are available as Suppl. Data File S2 and gene annotations for stramenopile CCRs as Suppl. Data File S3. **(B)** Residues involved in potassium selectivity in KCRs and the TM3 motif (105-109-116) across the subfamily of stramenopile CCRs and the related subfamilies of cryptophyte CCRs and uncharacterized proteins from Colpodellida.

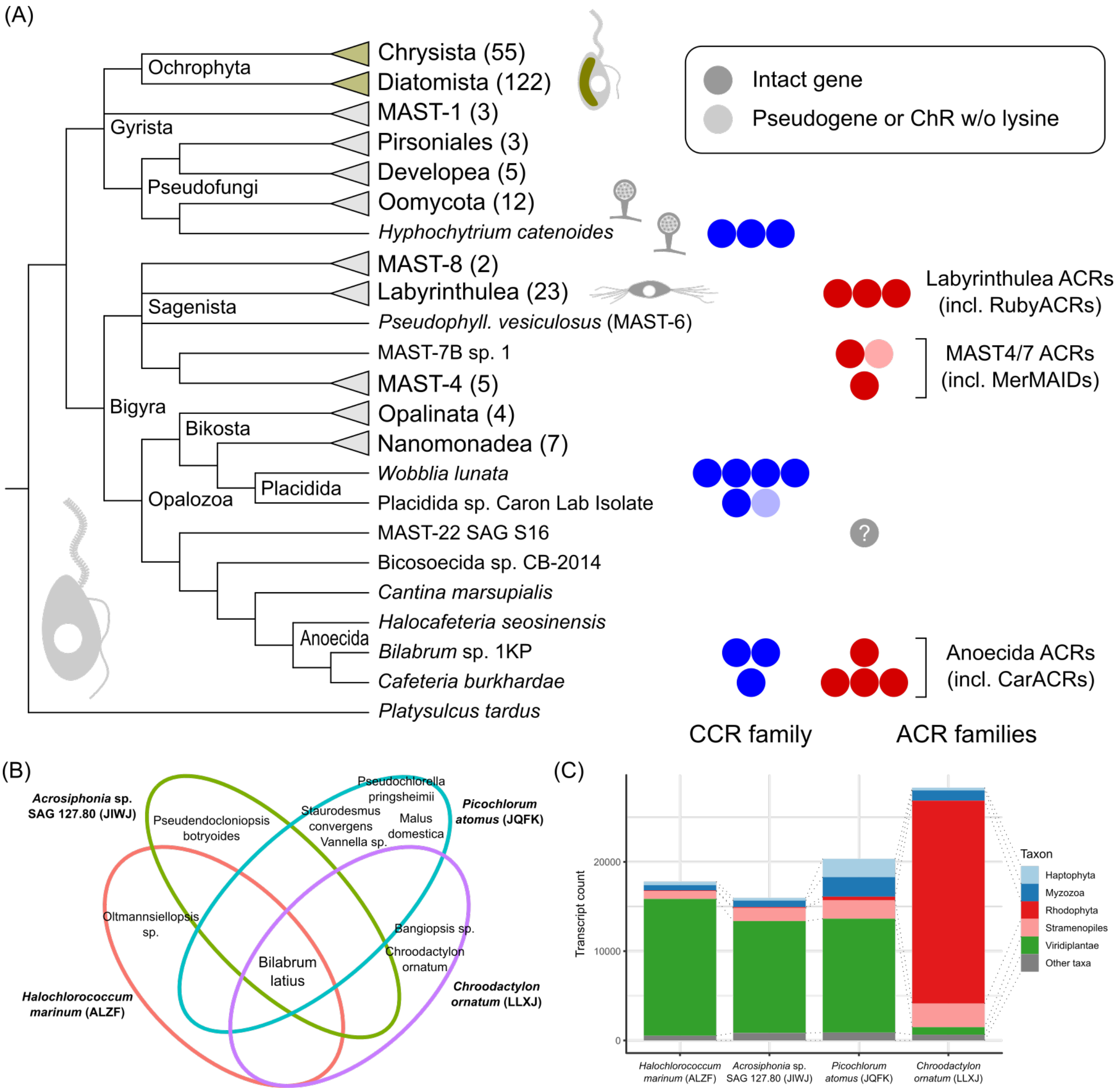


**Fig. S7. Stramenopile with ChR genes. (A)** Distribution of ChR families among Stramenopiles. Stramenopiles in the groundplan are heterotrophic heterokont flagellates. Appearance of fungus-like forms is indicated for hyphochytrids, oomycetes and labyrinthulids and of photosynthetic forms for ochrophytes. Blue and red dots correspond to individual ChR genes (note that the number of the ACR genes per assembly among Labyrinthulea varies). Numbers in parentheses indicate the number of species for which genetic data are available (for SAGs a comservative estimate is given). MAST (marine stramenopile) groups for which only single SAGs were available are omitted. Consensus cladogram according to (*6*–*9*). **(B)** and **(C)** ChRs B1ChR1, B1ChR2 and B1ChR3 appearing in transcriptome assemblies of four unrelated algae come from an anoecid stramenopile: **(B)** Incidence of SSU rRNA gene fragments in the four transcriptome assemblies. **(C)** Distribution of the best ublast hits among the protein sequences predicted for the four algae.

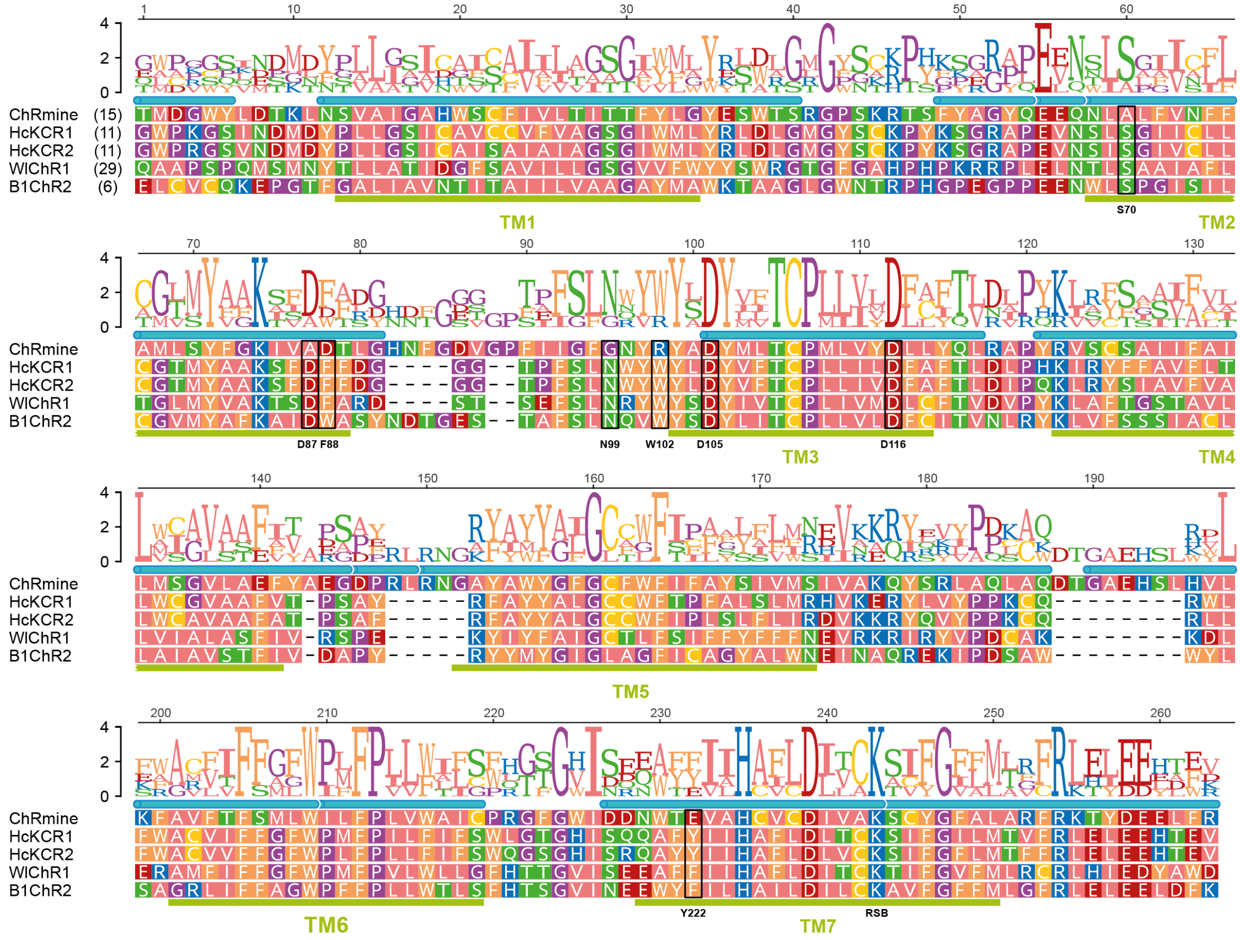


**Fig. S8**. **Structure-based multiple sequence alignment of KCRs and ChRmine** (see Suppl. Data File S4). Conservation for each position is represented as a sequence logo (y-axis reflecting bits of information). Secondary structure features are shown based on the structure of ChRmine (PDB: 7W9W): α-helical regions shown as cyan bars above and predicted transmembrane domains as green bars below. The alignment was trimmed at both ends to the rhodopsin domain proper with numbers in parentheses indicating the number of the residues trimmed from the N-terminus of each sequence.

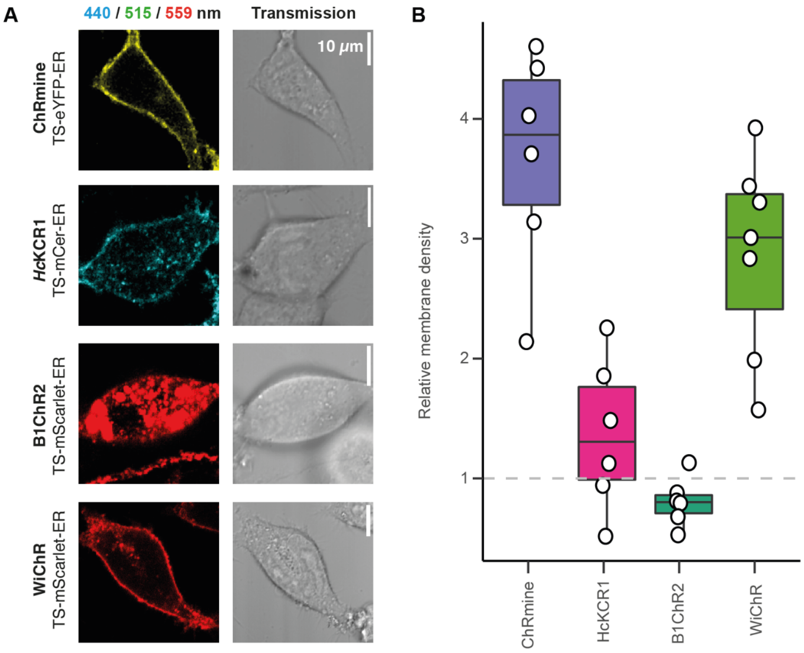


**Fig. S9. Membrane targeting of KCRs in ND7/23 cells (A)** Confocal images of ChRmine, and the potassium selective HcKCR1, WiChR, and B1ChR2 tagged with the named fluorescent proteins in ND7/23 cells. Scale bar: 10 µm. **(B)** Quantification of membrane targeting for the constructs visualized above. The targeting was calculated as the ratio between fluorescence density in the outer cell border and the enclosed intracellular fluorescence density.

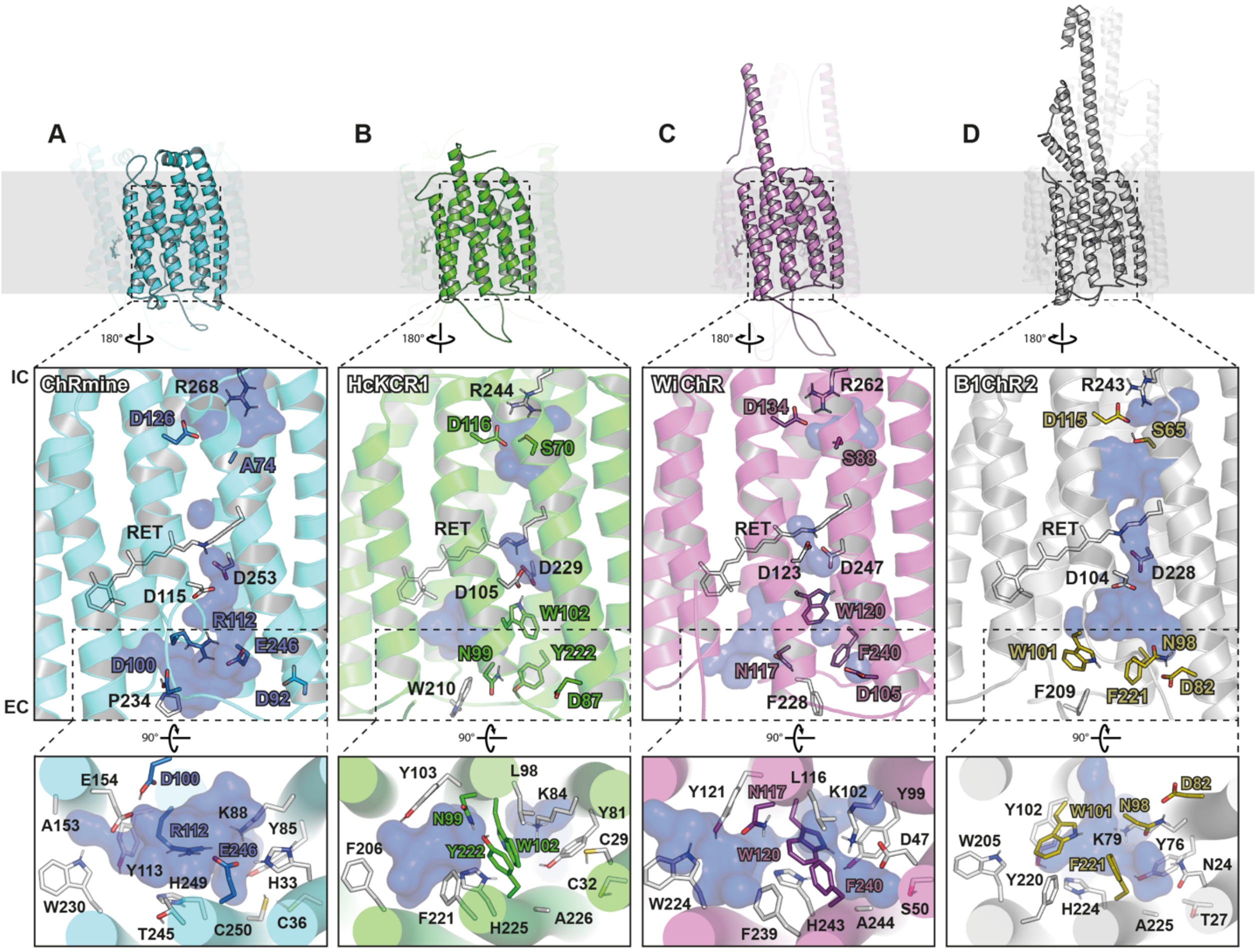


**Fig. S10**. Tertiary and 3D pore structures of **(A)** ChRmine based on an *in silico* equilibrated Cryo-EM structure (PDB-ID 7SFK) and $K^+$-selective BCCRs **(B)** HcKCR1, **(C)** WiChR and **(D)** B1ChR2 based on equilibrated AlphaFold2 models. Amino acids involved in cation selectivity and conduction in HcKCR1 and their homologues in other BCCRs are represented as licorice, those important for $K^+$-selectivity are highlighted in color. While selectivity filter amino acids in $K^+$-selective BCCRs are identical, their orientation in B1ChR2, especially N98 and W101, was found to be vastly different due to non-identical secondary residues. The same is true for counterion D104. Structurally, an overall high degree of conformity of HcKCR1 and WiChR, but not B1ChR2 was revealed. The KCR outer pore segments differ substantially from the ChRmine geometry, especially due to the abundance of aromatic amino acids resulting in different architectures of helices 2 and 3 and the extracellular loop 1. The model of B1ChR2 is an exception and its secondary structure has more similarity to ChRmine in that respect. Also, the pockets between helices 1, 2, 3 and 7 in WiChR and B1ChR2 are more polar because HcKCR1-C29 is exchanged to WiChR-D47 and B1ChR2-N24, thus attracting water molecules.

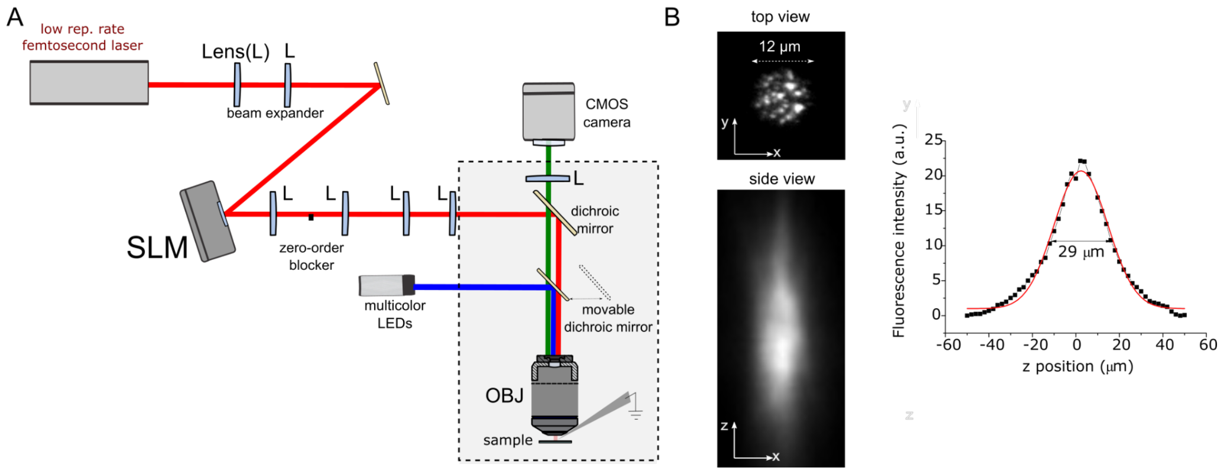


**Fig. S11.** Two-photon holographic stimulation system. **(A)** Schematic of the microscope. The holographic stimulation is based on the use of a low (500 kHz) repetition rate femtosecond laser at λ=1030 nm. The laser beam is phase-modulated by a liquid crystal based Spatial Light Modulator (SLM) and then projected at the pupil of the microscope objective (OBJ). Arbitrary illumination shape can be reproduced at the sample plane. **(B)** Top and side view of the fluorescence generated by a circular holographic spot (12 µm diameter), typically used for single cell photostimulation. Top and side view are acquired with the upper and a lower inverted objective respectively (see Methods). On the right, the integrated fluorescence axial profile: experimental point (black) and Gaussian fit (red) with an axial FWHM of 29 µm.

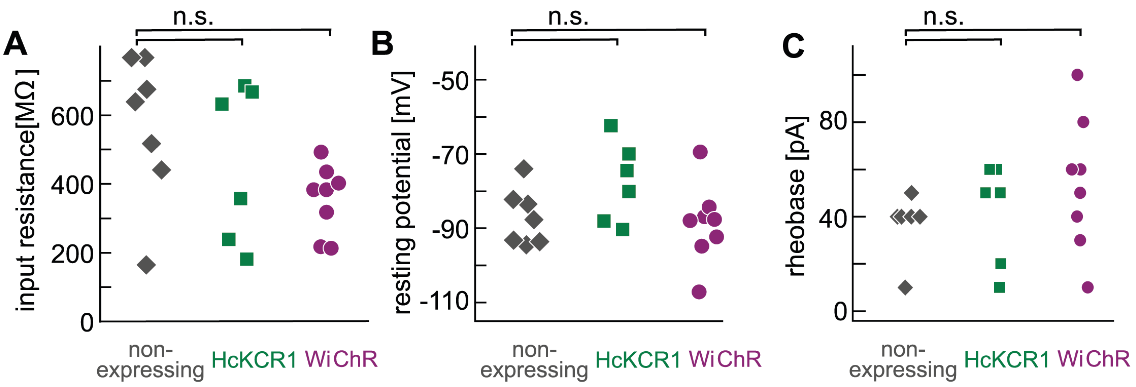


**Fig. S12**. **Neuronal membrane parameters** as determined right after establishing whole-cell configuration and switching to current-clamp mode for non-expressing neurons, neurons expressing HcKCR1 and neurons expressing WiChR. At a significance level of 0.01 (two-sample t-test) no differences were observed between expressing and non-expressing cells for all parameters. **(A)** Input resistance determined by injection of hyperpolarizing and depolarizing square currents (10 pA steps) and linear interpolation from the ΔV vs. ΔI curve. **(B)** Resting membrane potential determined within the first 30 s after break in. **(C)** Rheobase extracted as the first square current injection (1 s, 10 pA steps) that evoked an action potential.

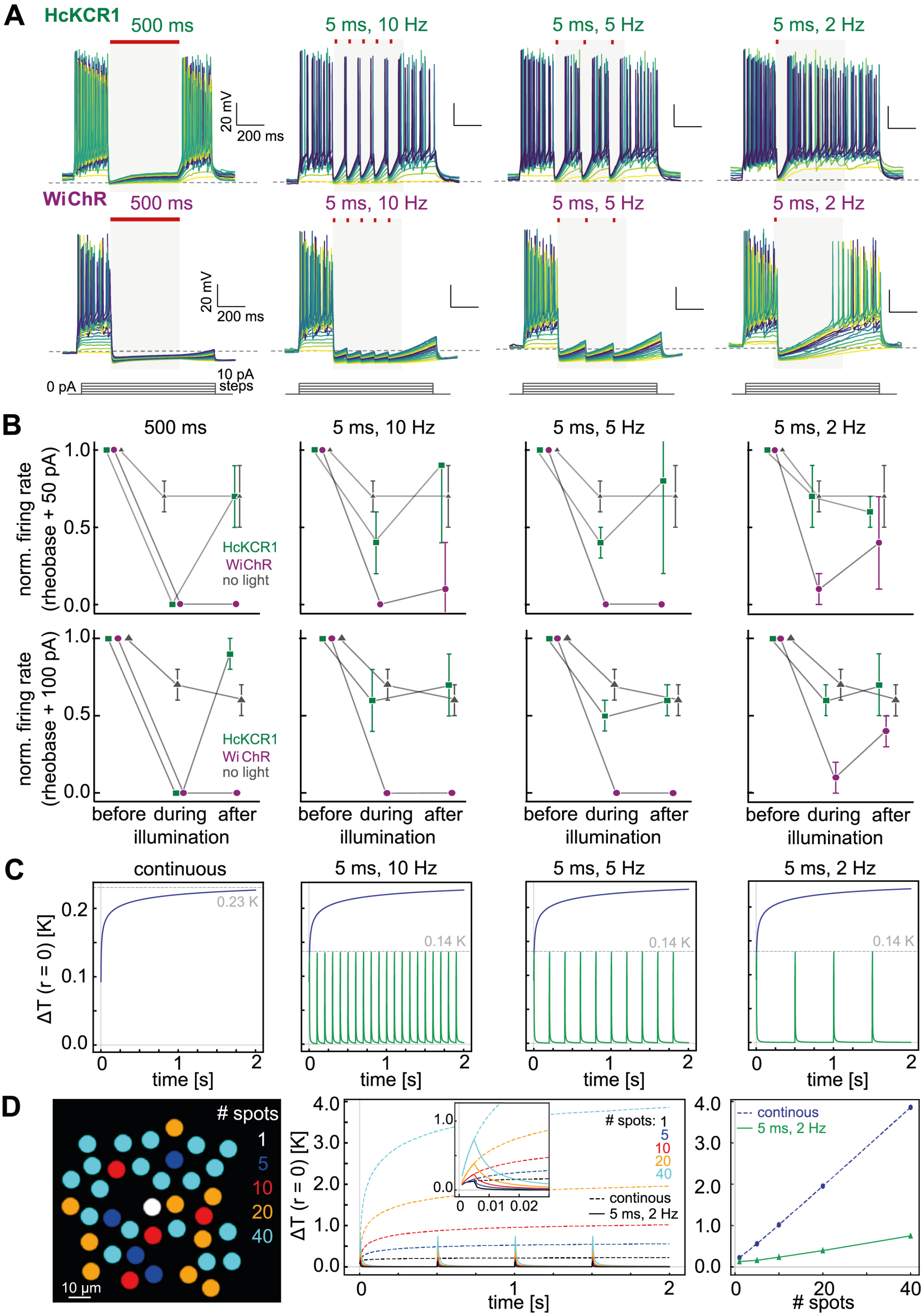
A
HcKCR1
500 ms
5 ms, 10 Hz
5 ms, 5 Hz
5 ms, 2 Hz
20 mV
200 ms
WiChR
0 pA
10 pA steps
B
norm. firing rate (rheobase + 50 pA)
norm. firing rate (rheobase + 100 pA)
HcKCR1
WiChR
no light
before during after
illumination
C
continuous
ΔT (r = 0) [K]
0.23 K
0.14 K
time [s]
D
### spots
10 µm
continous
5 ms, 2 Hz
### spots

206

**Fig. S13**. **Inhibition with pulsed 2-photon holographic illumination and corresponding local temperature rise in the tissue. (A)** Whole-cell current clamp recordings of HcKCR1 (top) and WiChR (bottom) with square current injections (1 s) of rising amplitude and an inhibition period of 500 ms in the middle. From left to right the duration of illumination is reduced from continuous for the whole 500 ms over 10 Hz and 5 Hz to 2 Hz pulsing of 5 ms illumination. All illumination protocols were compared at the same cell; power densities were the same for HcKCR1 and WiChR and the different frequencies (28-33 $\mu W/\mu m^2$) **(B)** Extraction of the firing rates for all pulsed protocols (and non-illuminated neurons) before (250 ms), during (500 ms) and after (250 ms) the illumination period normalized to the first 250 ms. Firing rates are shown for current injection 50 pA (left) and 100 pA (right) above the respective rheobase; determined as the first spike during the first or last 250 ms without light. Values are shown as mean±SD (HcKCR1_50pA n=6, WiChR_50pA n=7, noLight_50pA n=8, HcKCR1_100pA n=4, WiChR_50pA n=3, noLight_50pA n=3). **(C)** Simulation of local heating (r = 0) induced by one holographic spot (12 µm diameter, 31 $\mu W/\mu m^2$, 1030 nm) for continuous illumination (red) compared to pulsed 5 ms illumination (green) at different frequencies. Dashed lines indicate the peak temperature rise within 2 s **(D)** Simulation of local temperature rise for r = 0 (center of the first spot; white) as in (C) with increasing number of holographic spots placed within a 100 x 100 µm square (left); white - spot 1; blue - spots 2 - 5; red - spots 6 - 10; orange - spots 11 - 20; light blue - spots 21 - 40. Middle: time course of temperature rise for continuous (dashed line) *vs.* 2 Hz (cont. line) illumination for 1, 5, 10, 20 and 40 spots placed as shown on the left; inset shows the first 30 ms. Right: Extraction of the peak temperature rise after 2 s for indicated number of holographic spots comparing continuous (blue) and pulsed illumination (green).
